# Bioenergetic and Proteomic Profiling of ARID1A-Deficient Ovarian Clear Cell Carcinoma under Metabolic Challenge

**DOI:** 10.64898/2026.10.09.757928

**Authors:** Harith A. Ismail, Kristina Dankova, Trudy J. Philips, Brittany R. Davidson, Carly A. I. Twigg, Karthik Selvaraj, Valentino Clemente, Giancarlo Curci, Beatrice Dallan, Mihir Shetty, Asumi Hoshino, Pádraig D’Arcy, Francesco Nicoli, Shijia Zhu, Stefani N. Thomas, Martina Bazzaro

## Abstract

ARID1A loss is associated with increased mitochondrial respiration in ovarian clear cell carcinoma (OCCC), but whether this phenotype is maintained under metabolic challenge remains unclear. Here, we examined mitochondrial function and electron transport chain (ETC) protein abundance in isogenic ARID1A wild-type (WT) and ARID1A-knockout (KO) OCCC cells cultured under physiological glucose, galactose, or the Complex I inhibitor IACS-010759. Across all conditions, KO cells maintained higher basal, ATP-linked, and maximal respiration, greater spare respiratory capacity, and higher coupling efficiency than WT cells, despite lower steady-state ATP levels. In human OCCC tumors, ARID1A-negative cases showed a higher bioenergetic cellular (BEC) index than ARID1A-positive cases, and high BEC was associated with poorer cancer-specific survival. Quantitative proteomics revealed a trend toward higher ETC complex abundance in KO cells, although whole-complex differences did not reach statistical significance. ARID1A loss was associated with selective increases in Complex III and V proteins, whereas metabolic challenge was associated with increases in Complex I and IV proteins. Together, these findings show that the ARID1A-associated mitochondrial phenotype persists across distinct metabolic challenges and is not explained simply by coordinated increases in ETC protein abundance. The association of mitochondrial bioenergetics with clinical outcomes further supports its relevance in human OCCC.

**GRAPHICAL ABSTRACT:** 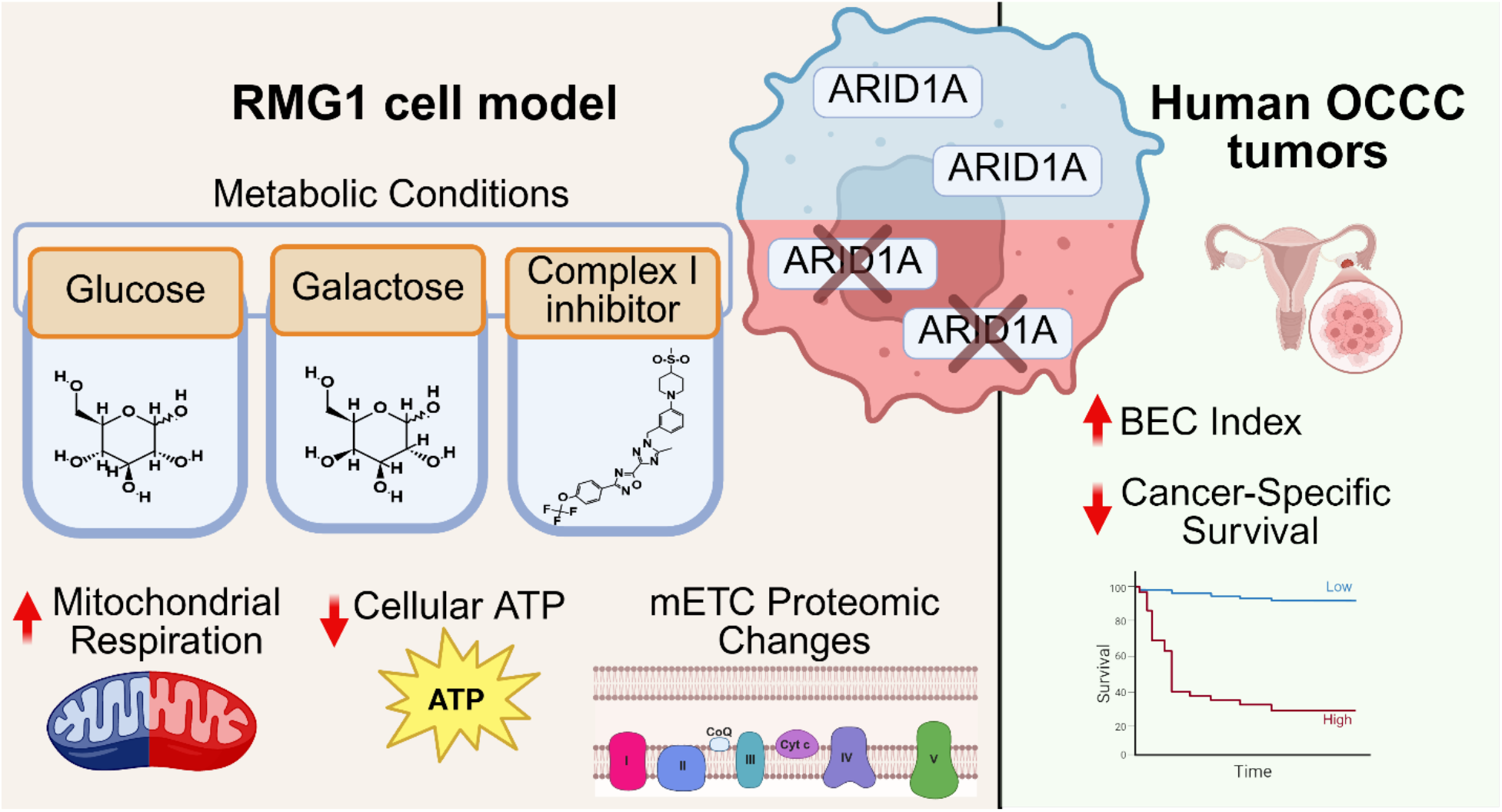

## INTRODUCTION

Ovarian clear cell carcinoma (OCCC) is a clinically and biologically distinct subtype of epithelial ovarian cancer characterized by poor response to platinum-based chemotherapy and poor prognosis in advanced-stage disease ^1–3^. Inactivating mutations in AT-rich interaction domain 1A (ARID1A), a core subunit of the switch/sucrose non-fermentable SWI/SNF chromatin-remodeling complex, occur in approximately 50–70% of OCCC cases ^4–6^. Beyond its canonical role in chromatin regulation, increasing evidence suggests that SWI/SNF complexes contribute to the regulation of cellular metabolism and mitochondrial function ^7^. Consistent with this, loss of ARID1A has been associated with metabolic rewiring and altered oxidative phosphorylation (OXPHOS) across multiple cancer contexts ^8,9^.

We have previously shown that ARID1A loss is associated with increased mitochondrial respiration in OCCC cells^10^. ARID1A depletion increased oxygen consumption, ATP-linked respiration, and maximal respiratory capacity, indicating increased reliance on oxidative metabolism. More recently, we demonstrated that ARID1A loss is also associated with remodeling of the mitochondrial proteome, including increased abundance of proteins involved in mitochondrial electron transport, mitochondrial translation, and ATP synthesis ^11^. These findings have highlighted mitochondrial metabolism as a potential vulnerability in ARID1A-deficient OCCC. However, whether this mitochondrial phenotype is maintained when cells encounter different metabolic demands remains unclear.

Metabolic dependencies are not static and can be influenced by nutrient availability and compensatory responses to respiratory inhibition ^12,13^. This is particularly relevant for OCCC, in which glucose metabolism is a prominent feature of tumor biology, and glucose availability has been shown to influence tumor cell survival under metabolic stress ^14,15^. Because such adaptations may alter the effectiveness of mitochondria-targeted strategies, determining whether ARID1A-associated mitochondrial dependence is maintained or reshaped under metabolic challenge is important for identifying vulnerabilities that remain therapeutically actionable.

To address this question, we used two complementary approaches to challenge mitochondrial dependence. Replacing glucose with galactose reduces reliance on glycolytic ATP production and increases dependence on mitochondrial oxidative phosphorylation for maintenance of cellular energy homeostasis ^16–18^. As a complementary pharmacological approach, we used IACS-010759, a potent and selective inhibitor of mitochondrial electron transport chain (ETC) Complex I that was developed to target OXPHOS-dependent cancer cells ^19,20^. IACS-010759 directly suppresses mitochondrial respiration, providing a defined means of challenging OXPHOS-dependent metabolism ^19^. Together, these perturbations allow us to determine whether the mitochondrial phenotype associated with ARID1A loss is maintained under conditions that increase mitochondrial dependence or directly challenge respiratory function.

In this study, we investigated how ARID1A-deficient OCCC cells respond to defined metabolic challenges using mitochondrial respiratory analysis, cellular ATP measurements, and quantitative proteomics. RMG1 parental and ARID1A-knockout (D9) cells were exposed to control medium, galactose-based medium, or pharmacological inhibition of mitochondrial Complex I with IACS-010759. Mitochondrial respiratory function was assessed using Seahorse XF metabolic flux analysis, while quantitative proteomic mass spectrometry-based profiling was used to characterize associated changes in protein abundance and biological pathways. In a separate clinical analysis, we examined the bioenergetic cellular (BEC) index in human OCCC specimens according to ARID1A status and evaluated its association with cancer-specific survival.

## EXPERIMENTAL PROCEDURES

### Cell Culture and Metabolic Perturbations

RMG1 wild-type (WT) and ARID1A-knockout (D9) ovarian clear cell carcinoma cells were obtained from Dr. Pádraig D’Arcy (Linköping University, Sweden). The D9 clone was previously generated and characterized alongside an independent ARID1A-knockout RMG1 clone (G11); both clones showed efficient ARID1A depletion, and D9 was selected for downstream analyses because of its greater depletion efficiency ^11^. Cells were maintained in high-glucose Dulbecco’s Modified Eagle’s Medium (DMEM) GenClone # 25-2500 containing sodium pyruvate, L-glutamine, and phenol red and supplemented with 10% fetal bovine serum (FBS). Cells were cultured at 37°C in a humidified incubator containing 5% CO₂.

For metabolic perturbation experiments, cells were cultured under three conditions: (1) control medium consisting of DMEM containing 1 g/L glucose (∼5.5 mM), sodium pyruvate, L-glutamine, and phenol red, supplemented with 10% FBS, reflecting physiological tumor glucose concentrations (hereafter, Control); (2) galactose medium consisting of phenol red-free, glucose-free DMEM supplemented with 10 mM galactose, 6 mM L-glutamine, 5 mM HEPES, 1 mM sodium pyruvate, and 10% FBS (hereafter Galactose or Gal); or (3) control medium supplemented with 10 nM IACS-010759, a mitochondrial electron transport chain Complex I inhibitor (Selleck Chemicals, Houston, TX, USA; cat. no. S8731) (hereafter, Complex I inhibitor or C1). Cells were exposed to the indicated conditions for 16 h prior to downstream analysis.

### Cellular ATP Measurement

Cellular ATP levels were measured using the CellTiter-Glo Luminescent Cell Viability Assay following 16 h exposure to the indicated metabolic conditions. Equal numbers of RMG1 WT and ARID1A-knockout D9 cells were seeded for each experimental condition. ATP concentrations were determined using an ATP standard curve. Experiments were performed in three independent biological replicates, with technical triplicates within each experiment.

### Seahorse XF Cell Mito Stress Test

Mitochondrial respiration was assessed using the Seahorse XF Cell Mito Stress Test Kit on a Seahorse XF96 Analyzer (Agilent Technologies) according to the manufacturer’s instructions. RMG1 WT and ARID1A-knockout D9 cells were seeded in XF96 cell culture microplates at 40,000 cells per well, with 12 wells per clone per experimental condition, and allowed to adhere overnight. Cells were subsequently exposed to the indicated metabolic conditions prior to Seahorse analysis. Immediately prior to measurement, the treatment medium was replaced with Seahorse XF assay medium containing 10 mM glucose, 2 mM glutamine, 1 mM pyruvate, and 5 mM HEPES, adjusted to pH 7.4. Plates were equilibrated for 1 h at 37°C in a non-CO₂ incubator. Oligomycin (1.5 µM), FCCP (2 µM), and rotenone/antimycin A (0.5 µM) were sequentially injected during the assay. Oxygen consumption rate (OCR) measurements were used to determine basal respiration, ATP-linked respiration, maximal respiration, spare respiratory capacity, proton leak, and coupling efficiency. Parameters were calculated using Wave software (Agilent Technologies, version 2.6). Basal respiration was calculated as baseline OCR minus non-mitochondrial OCR; ATP-linked respiration as baseline OCR minus the minimum OCR following oligomycin injection; maximal respiration as the maximum OCR following FCCP injection minus non-mitochondrial OCR; and spare respiratory capacity as maximal respiration minus basal respiration. Coupling efficiency and percentage spare respiratory capacity were expressed relative to basal respiration. Wells failing technical quality-control criteria or exhibiting z-scores >2.5 were excluded from analysis. Statistical comparisons were performed using ordinary two-way ANOVA with cell genotype and metabolic condition as fixed factors, followed by Šídák-adjusted multiple comparisons between RMG1 WT and ARID1A-knockout D9 cells within each metabolic condition. Analyses were performed using GraphPad Prism version 11.1.0. Data are presented as mean ± SEM, and adjusted p < 0.05 was considered statistically significant.

### Ethics statement

Previously collected OCCC tissue specimens were used with approval from the Institutional Review Board of the University of Minnesota (STUDY00006529). All tissues were obtained with written informed consent from the patients, and the study was conducted in accordance with the ethical principles of the Belmont Report.

### Immunohistochemistry analysis

Formalin-fixed, paraffin-embedded (FFPE) OCCC specimens from a previously characterized patient cohort were analyzed by immunohistochemistry. Representative tumor cores were arranged in tissue microarrays, and 5-µm sections were deparaffinized and rehydrated by sequential washing with xylene, 100% ethanol, 95% ethanol, 80% ethanol, and PBS. Antigen retrieval was performed using 1× Reveal Decloaker (Biocare Medical) for 30 min at 100°C, followed by blocking with Background Sniper (Biocare Medical) for 13 min at room temperature. Sections were incubated with antibodies against ARID1A, TOM20, or ATP5B. Immunoreactivity was visualized using 3,3′-diaminobenzidine and sections were counterstained with hematoxylin.Immunohistochemical staining was quantified using an H-score, calculated by multiplying the staining intensity score (0 = no staining, 1 = weak staining, 2 = moderate staining, and 3 = strong staining) by the corresponding percentage of positively stained cells, resulting in a score ranging from 0 to 300. The bioenergetic cellular (BEC) index was calculated as the ratio of the ATP5B H-score to the TOM20 H-score. A total of 53 OCCC specimens were included in the BEC analysis, comprising 35 ARID1A-positive and 18 ARID1A-negative tumors.

### Protein Extraction and Sample Preparation for Proteomic Analysis

RMG1 WT and ARID1A-knockout D9 cells were exposed to control, galactose, or Complex I inhibitor conditions for 16 h prior to sample collection, with three biological replicates per group. Following treatment, cells were placed on ice, washed with ice-cold phosphate-buffered saline (PBS), and harvested by scraping in cold PBS. Cell pellets were collected by centrifugation at 300–400 × g for 5 min at 4°C and stored at −80°C until further processing. Cell pellets (n=3 biological replicates) were resuspended in 50µl lysis buffer (8 M urea, 75 mM NaCl, 50 mM Tris-HCl (pH 8.0), 1 mM EDTA, 1mM PMSF, 2µg/mL aprotonin, 10µg/mL leupeptin), vortexed for 10 s, incubated on ice for 15 min and repeated. Lysates were clarified by centrifugation at 17,000 × g for 10 min at 4°C, supernatant was transferred to a new 1.5 mL tube. Protein concentrations were determined using the Bicinchoninic Assay (Thermo Fisher Scientific). A pooled sample containing an equivalent amount of protein from all samples was prepared to facilitate inter-batch normalization. For enzymatic digestion, 35 µg of protein from each sample was reduced with 5 mM dithiothreitol for 1 h at 37°C and alkylated with 10 mM iodoacetamide for 45 min at 25°C in the dark. Samples were diluted 1:3.5 (v/v) with 50 mM Tris-HCl (pH 8.0) and subjected to sequential enzymatic digestion, first with LysC (Wako) at a ratio of 1 mAU per 50 µg protein for 2 h at 25°C, followed by trypsin (Promega) at a 1:50 enzyme-to-protein ratio (w/w) for 18 h at 25°C. Digestion was terminated by acidification with formic acid to a final concentration of 1% (v/v). Peptides were desalted using 1-mg C18 Sep-Pak cartridges (Waters) and concentrated by vacuum centrifugation.

### Tandem Mass Tag Labeling

Dried peptides were reconstituted in 40 µL of 50 mM HEPES (pH 8.5) by sonication for 3 min. Insoluble material was removed by centrifugation at 13,000 rpm for 3 min at room temperature. Equal amounts of peptide from each sample were labeled using TMT 10-plex Mass Tag Labeling Reagents (Thermo Fisher Scientific) according to the manufacturer’s protocol, using a TMT-to-peptide ratio of 7.5:1 (w/w) and a TMT reagent concentration of 20 µg/µL. The TMT labeling scheme was as follows: Plex 1 – WT LG_3 (126), D9 Gal_2 (127N), WT C1_1 (127C), D9 Gal_1 (128N), WT C1_2 (128C), D9 C1_1 (129N), WT Gal_1 (129C), WT LG_1 (130N), D9 LG_3 (130C); Plex 2 – WT LG_2 (126), D9 C1_3 (127N), D9 Gal_3 (127C), WT C1_3 (128N), WT Gal_3 (128C), D9 LG_1 (129N), D9 LG_2 (129C), WT Gal_2 (130N), D9 C1_2 (130C); Pool (131). Labeling efficiency was verified to exceed 97% before quenching with 0.2% hydroxylamine (v/v). Labeled samples were combined, desalted using C18 STAGE Tips, and concentrated by vacuum evaporation.

### Basic Reversed-Phase Liquid Chromatography

Dried TMT-labeled peptides were reconstituted in 2% acetonitrile containing 4.5 mM ammonium formate by sonication for 3 min. Insoluble material was removed by centrifugation at 13,000 rpm for 3 min at room temperature. Peptide fractionation was performed by basic reversed-phase liquid chromatography using an Agilent 1200 Infinity II system equipped with a ZORBAX 300 Extend-C18 column (100 mm × 2.1 mm, 3.5 µm particle size; Agilent Technologies, Santa Clara, CA, USA) at a flow rate of 200 µL/min. Mobile phase A consisted of 2% acetonitrile and 4.5 mM ammonium formate in water, whereas mobile phase B consisted of 90% acetonitrile and 4.5 mM ammonium formate. Peptides were fractionated using a linear gradient of 0–16% B over 6 min, 16–40% B over 60 min, 40–44% B over 4 min, and 44–60% B over 5 min. Fractions were collected at 1-min intervals and concatenated into 18 final fractions. Samples were frozen at −80°C for 30 min and concentrated by vacuum evaporation. Prior to LC-MS/MS analysis, peptides were quantified using a NanoDrop spectrophotometer (Thermo Fisher Scientific).

### Liquid Chromatography–Tandem Mass Spectrometry Data Acquisition

LC-MS/MS analysis was performed using a Vanquish Horizon liquid chromatography system coupled with an Orbitrap Q Exactive Plus mass spectrometer (Thermo Fisher). Peptides (2 µg) were injected onto an Acclaim PepMap 100 C18 HPLC column (Thermo Scientific; 1 x 150mm; particle size: 2 µm). Peptides were separated at an analytical flow rate of 0.3 mL/min using a linear gradient between solvent A (0.1% formic acid in water) and solvent B (0.1% formic acid in 80% ACN) as follows: 5% Solvent B at 0–1.0 min, 5-40% Solvent B at 1.0–61.0 min, 40–95% Solvent B at 61.0–62.0 min, 95-5% Solvent B at 62.0–66.0 min, and 5% Solvent B at 66.0–69.0 min. Data were acquired using a top 10 DD-MS^2^ method. MS1 scans were acquired in the Orbitrap over a scan range of 200-2000 m/z at a resolution of 35,000 with a 4-second dynamic exclusion duration. MS2 scans were acquired with an isolation window of 2.0 m/z. The raw mass spectrometry data were deposited in the ProteomeXchange Consortium through the PRIDE repository under accession PXD084774.

### Mass Spectrometry Data Analysis

The raw MS data were processed using Proteome Discoverer 3.0 (Thermo Fisher Scientific) with CHIMERYS (Thermo Fisher Scientific) for protein identification using the following parameters: enzyme specificity: trypsin; maximum missed cleavage sites: 2; peptide (MS1) and fragment mass tolerance: 20 ppm and 0.6 Da, respectively. Oxidation of methionine and acetylation of the protein N-terminus were set as variable modifications, and cysteine carbamidomethylation was set as a fixed modification. Percolator was used to determine the confidence for peptide and protein identification; validation was based on a q-value < 0.05, and a strict false discovery rate (FDR) target was set to 0.01. For TMT-labeled protein quantification, reporter ion MS2 was selected, and isobaric labels with their correction factors were applied from a TMT 10-plex and multiplexed into two sets. All other parameters within PD remained under default settings. For protein and peptide quantification, the peak integration tolerance of ‘Reporter Ions Quantifier’ was set to 20 ppm. Spectra were searched against a UniProt/Swiss-Prot *Homo sapiens* protein database (downloaded 19 Sept. 2026) together with common contaminant sequences. Proteins were filtered based on high protein FDR confidence, contaminant set to false, and abundances greater than 0 in every sample. Following data filtering, data were exported to Excel. Protein intensities, including the pooled reference samples, were first median-normalized to correct for differences in overall signal intensity. To account for variation between the two sets (analytical runs), the median-normalized intensities were scaled using an internal reference scaling factor (IRS) calculated from the corresponding pooled-referenced sample. Data were transferred to Perseus (v 2.6.1) for analysis. Data were log_2_-transformed, and the non-valid values were removed. The residual inter-run batch effects were corrected in Perseus using the “removeBatchEffect” function from the limma R package done through the Perseus R plugin. Differential protein abundances for the different comparisons were assessed using an FDR of 0.01 and an s0 factor estimated using the fudge2 function in R.

Protein-protein interaction (PPI) networks were generated for the top N upregulated or downregulated proteins driving pathway enrichment using STRING-db ver 12.0 (string-db.org) and filtered for high-confidence interactions (0.900).

### Statistical Analysis

For Seahorse experiments, statistical comparisons were performed using ordinary two-way ANOVA with cell genotype and metabolic condition as fixed factors, followed by Šídák-adjusted multiple comparisons between RMG1 WT and ARID1A-knockout D9 cells within each metabolic condition. Analyses were performed using GraphPad Prism version 11.1.0. Data are presented as mean ± SEM, and adjusted p < 0.05 was considered statistically significant.

For cellular ATP measurements presented in Figure 1D, statistical comparisons between RMG1 WT and D9 cells within each metabolic condition were performed using an unpaired two-tailed t-test. Data are presented as mean ± SEM of technical triplicates, and *p* < 0.05 was considered statistically significant.

**Figure 1.**
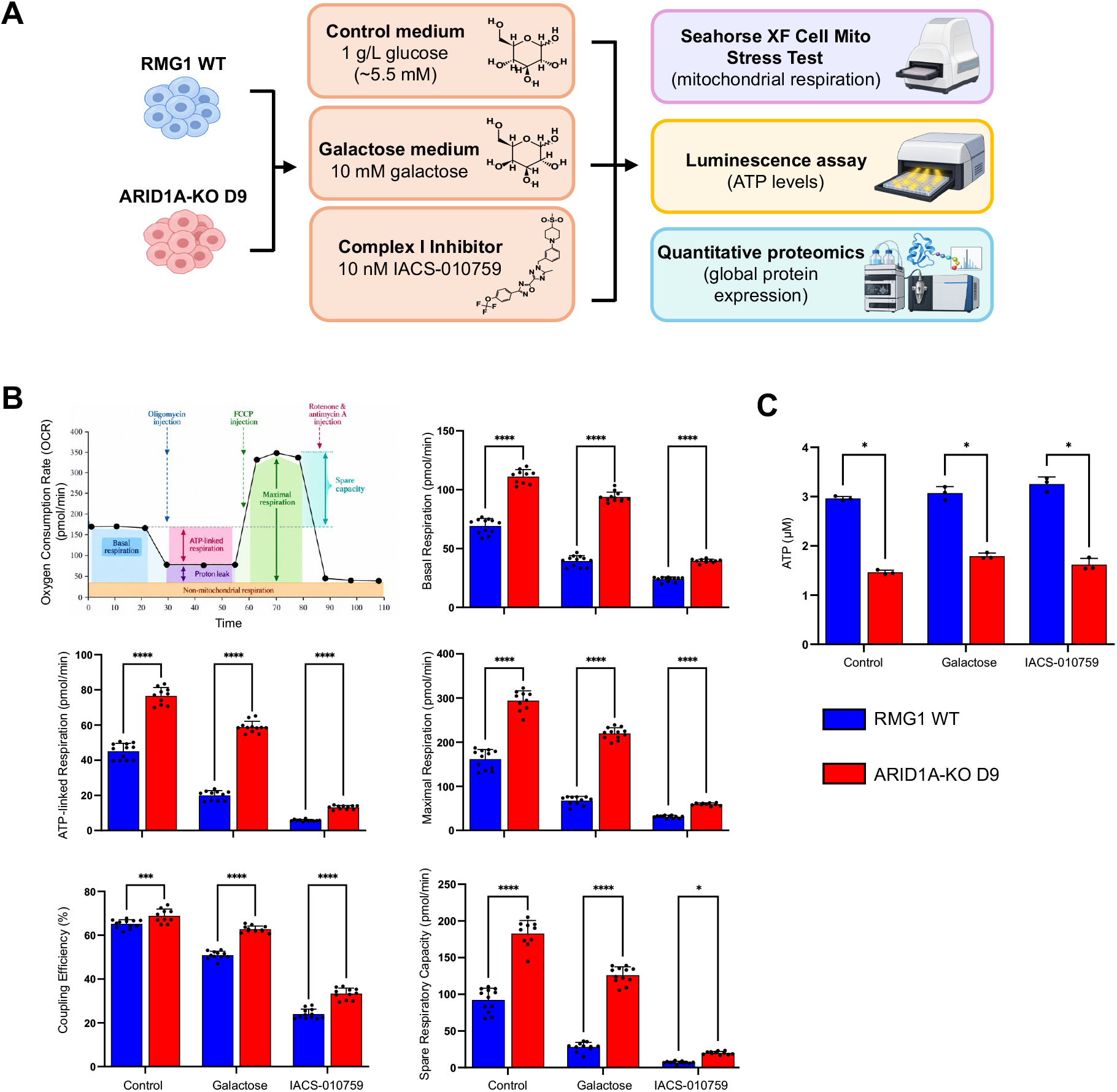
ARID1A loss is associated with increased mitochondrial respiration and reduced ATP levels under metabolic challenge. **(A)** Experimental design. RMG1 wild-type (WT) and ARID1A-knockout D9 cells were exposed for 16 h to control medium containing 1 g/L glucose (∼5.5 mM), galactose medium containing 10 mM galactose, or control medium supplemented with 10 nM IACS-010759. Cells were subsequently analyzed by Seahorse XF Cell Mito Stress Test, cellular ATP measurement, and quantitative proteomics. **(B)** Mitochondrial respiratory parameters measured using the Seahorse XF Cell Mito Stress Test. The schematic illustrates respiratory responses following sequential addition of oligomycin, FCCP, and rotenone/antimycin A. Basal respiration, ATP-linked respiration, maximal respiration, coupling efficiency, and spare respiratory capacity are shown for WT and D9 cells under the indicated conditions. Individual points represent analyzed Seahorse wells, and bars represent mean ± SEM. Statistical comparisons between WT and D9 cells within each metabolic condition were performed using ordinary two-way ANOVA followed by Šídák-adjusted multiple comparisons. *p < 0.05, ***p < 0.001, ****p < 0.0001. **(C)** Cellular ATP concentrations in RMG1 WT and ARID1A-knockout D9 cells following 16 h exposure to the indicated metabolic conditions. ATP concentrations were determined using an ATP standard curve. Bars represent mean ± SEM of technical triplicates. Statistical comparisons between WT and D9 cells within each metabolic condition were performed using an unpaired two-tailed t-test.

For the BEC analysis, differences between ARID1A-positive and ARID1A-negative tumors were assessed using an unpaired two-tailed Welch’s t-test, and cancer-specific survival was evaluated using Kaplan–Meier analysis.m The statistical analyses of the proteomic data included two-way ANOVA followed by Benjamini-Hochberg (BH) correction, and limma-moderated t-tests followed by BH correction.

## RESULTS

### ARID1A Loss Is Associated with Increased Mitochondrial Respiration and Reduced ATP Levels Under Metabolic Challenge

The RMG1 D9 clone used in this study was previously characterized alongside an independent ARID1A-deficient RMG1 clone (G11), with D9 selected for downstream analyses based on its greater ARID1A depletion efficiency ^11^. To determine whether the mitochondrial respiratory phenotype associated with ARID1A loss is maintained under different metabolic conditions, mitochondrial respiration was assessed in RMG1 WT and ARID1A-knockout D9 cells following exposure to control, galactose, or the Complex I inhibitor IACS-010759 (**Figure 1A**). Across all three conditions, D9 cells displayed higher basal respiration, ATP-linked respiration, and maximal respiration compared with WT cells (**Figure 1B**). D9 cells also exhibited greater spare respiratory capacity and higher coupling efficiency across the metabolic conditions examined (**Figure 1B**). Although respiratory activity decreased following Complex I inhibition in both cell types, D9 cells maintained higher respiratory parameters than WT cells. Additional Seahorse-derived parameters, including proton leak, non-mitochondrial respiration, and spare respiratory capacity expressed as a percentage of basal respiration, are shown in **Supplementary Figure S1A**.

Cellular ATP levels were also assessed following exposure to the same metabolic conditions. D9 cells exhibited lower ATP concentrations than WT cells under control, galactose, and Complex I inhibitory conditions (**Figure 1C**). These findings indicate that increased mitochondrial respiratory activity in D9 cells was accompanied by lower steady-state cellular ATP abundance under the conditions examined.

### ARID1A Loss Is Associated with an Increased Bioenergetic Cellular Index in OCCC Tumors

The bioenergetic cellular (BEC) index has been used to assess tumor bioenergetic state ^21^. To determine whether the mitochondrial phenotype associated with ARID1A loss is also evident in clinical OCCC specimens, we next examined the BEC index of 53 tumors from a previously characterized OCCC cohort ^22^. In this cohort, BEC was calculated as the ATP5B/TOM20 ratio from immunohistochemical staining, with ATP5B representing a subunit of mitochondrial ATP synthase and TOM20 serving as a marker of mitochondrial abundance. Representative ARID1A- positive (ARID1A+) and ARID1A-negative (ARID1A-) tumors with corresponding TOM20 and ATP5B staining are shown in **Figure 2A**. ARID1A-negative tumors exhibited significantly higher BEC index than ARID1A-positive tumors (**Figure 2B**), indicating increased ATP synthase abundance relative to mitochondrial content. Furthermore, patients with a high BEC index exhibited poorer cancer-specific survival compared with patients with a low BEC index (p = 0.0017; HR = 5.1, 95% CI 1.4–18; **Figure 2C**). Together, these findings extend the mitochondrial phenotype observed in ARID1A-deficient RMG1 cells to clinical OCCC specimens and suggest that a higher BEC index, as a proxy of mitochondrial bioenergetic activity, is associated with adverse clinical outcome.

**Figure 2.**
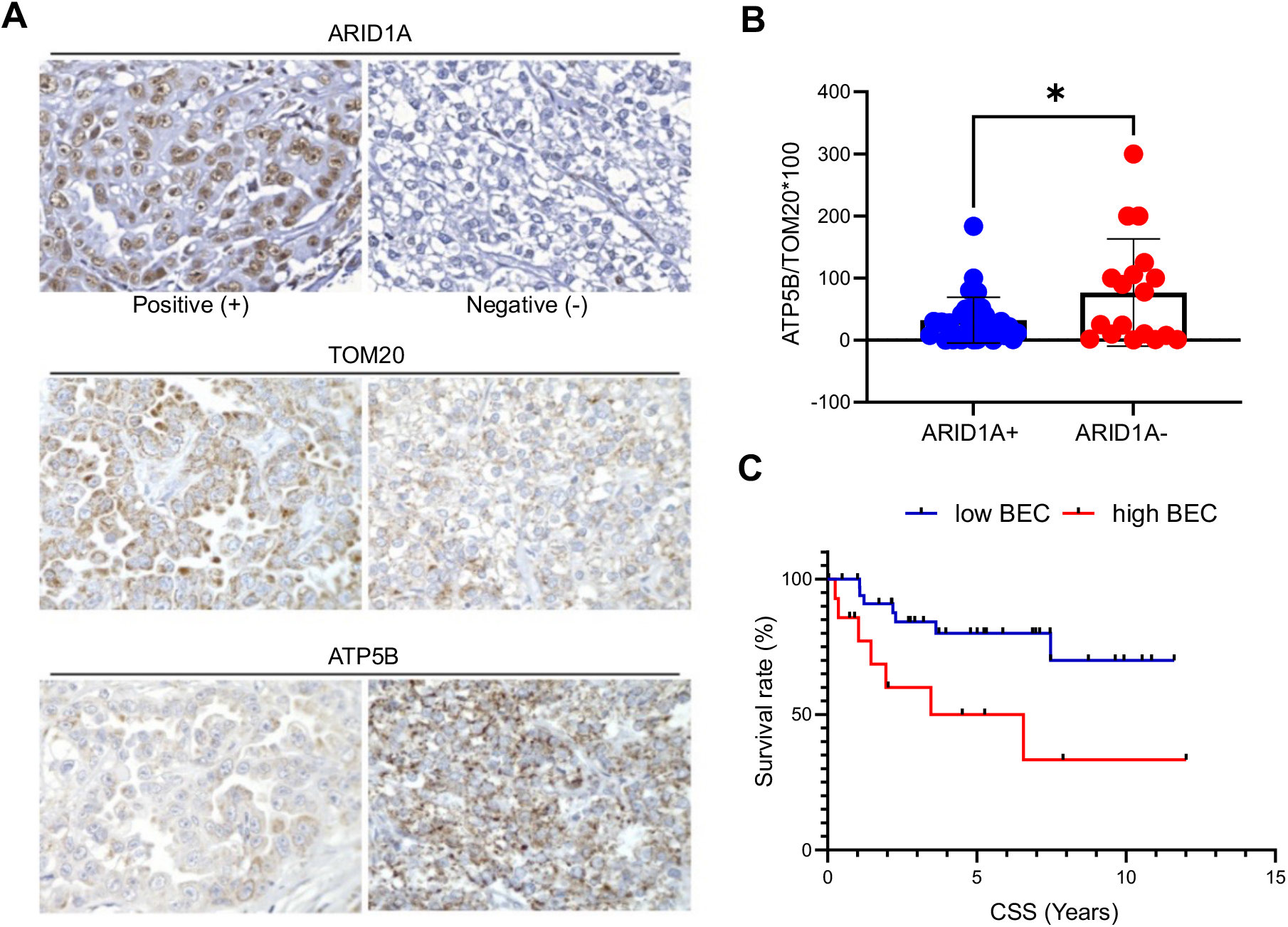
ARID1A deficiency is associated with a higher bioenergetic cellular index in human OCCC. **(A)** Representative immunohistochemical staining of ARID1A, TOM20, and ATP5B in ARID1A-positive (ARID1A+) and ARID1A-negative (ARID1A−) OCCC specimens. **(B)** BEC index, calculated as ATP5B/TOM20 H-score × 100, in ARID1A-positive (n = 35) and ARID1A-negative (n = 18) OCCC specimens. Individual points represent individual tumors. Statistical significance was determined using an unpaired two-tailed Welch’s t-test; p = 0.0498. **(C)** Kaplan-Meier analysis of cancer-specific survival according to BEC index. Patients were classified as low BEC (<0.5) or high BEC (≥0.5). High BEC was associated with poorer cancer-specific survival (p = 0.0017, HR = 5.1, 95% CI = 1.4–18).

### Proteomic Responses to Metabolic Challenge in ARID1A-Deficient OCCC Cells

To interrogate the perturbations associated with ARID1A loss and metabolic challenge in RMG1 cells, we conducted a TMT-based global proteomic analysis using analytical flow liquid chromatography. A total of 5203 proteins were identified, 4554 proteins were quantified, and 3868 proteins were quantified without any missing or imputed values across all samples (n = 18). Each of the six conditions – ARID1A WT and D9 cells: control, galactose, and Complex I inhibitor treatment – were analyzed in biological triplicate, with strong protein abundance correlations (r = 0.987 – 0.996) among the biological replicates (**Supplementary Figure S2** and **Table S1**).

Principal Component Analysis revealed distinct clustering of the WT vs. D9 proteomes (**Figure 3A**). Whereas the proteomes of the galactose-treated WT cells were highly similar among the biological replicates, galactose treatment of the D9 cells resulted in greater differences among the proteomes of the biological replicates. In both cell lines, treatment with the Complex I inhibitor IACS-010759 induced proteome changes that were more similar to the control condition than galactose treatment, as evidenced by the hierarchical clustering results (**Figure 3B**).

**Figure 3.**
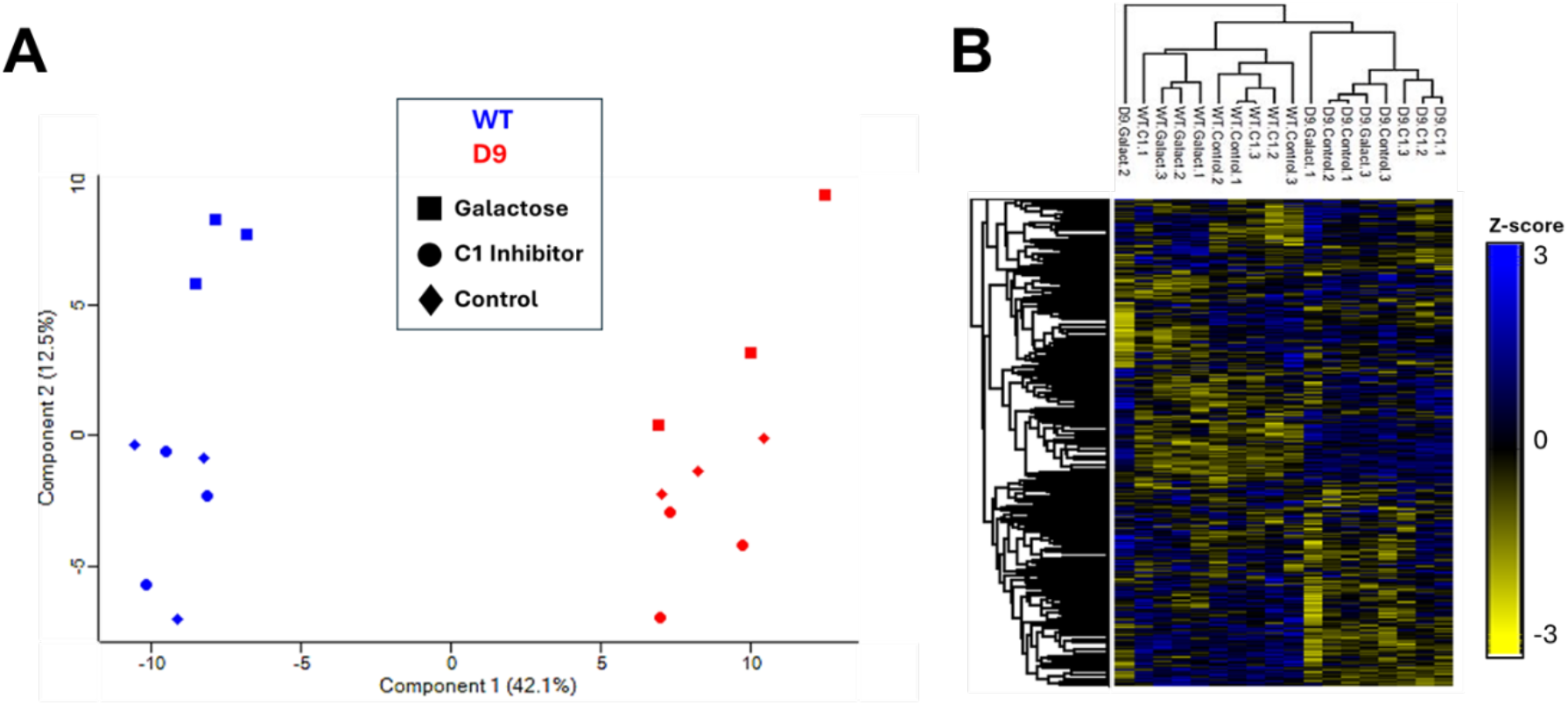
Clustering of proteomes and proteins associated with ARID1A loss and metabolic stress induced by galactose and inhibition of Complex I activity. **A)** Principal component analysis (PCA) plot. **B)** z-score-based hierarchical clustering of proteins quantified across all samples without any missing values (n = 3868).

Based on the number of differentially expressed proteins, treatment with the metabolic stressors altered the RMG1 proteome to a lesser extent compared to the loss of ARID1A expression. A total of 94 proteins had significant differences in relative abundance when comparing the D9 vs. WT proteomes in the control condition (**Figure 4A**). Pathway analysis of the proteins whose abundances increased following the loss of ARID1A expression indicated an enrichment of biological processes such as translation and cellular macromolecule biosynthesis (**Figure 4B**).

**Figure 4.**
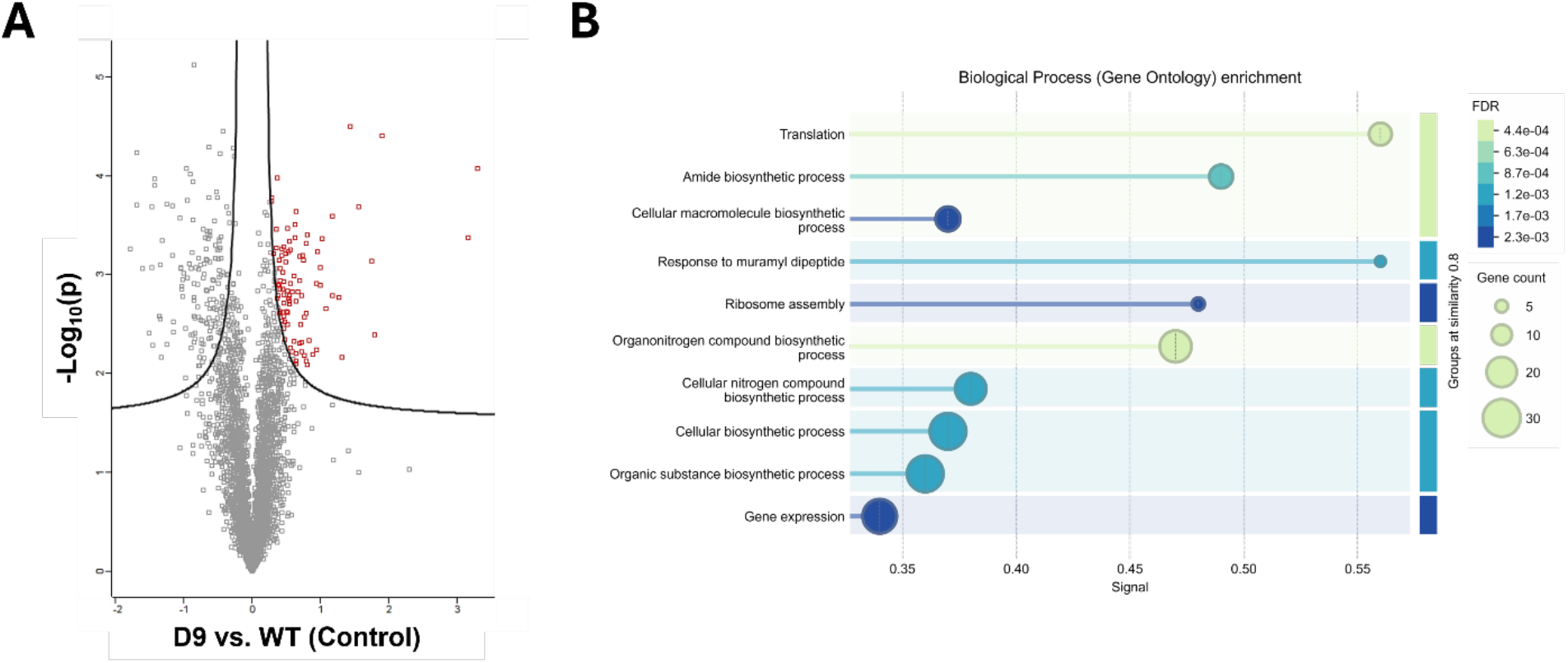
Loss of ARID1A expression is associated with global proteome perturbations. **A)** Differential protein expression among the ARID1A KO (D9) vs. WT RMG1 cells. Significantly dysregulated proteins are indicated in red (up-regulated proteins). Permutation-based false discovery rate (FDR, *p* < 0.01, s_0_ = 0.5206) estimation is visualized with black hyperbolic curves. **B)** Enriched Reactome Biological Process pathways among the up-regulated proteins associated with ARID1A loss.

Of particular interest with the global proteomic analysis of the RMG1 cells was insight into how the mitochondrial ETC protein abundances would be altered in response to the metabolic challenges described in **Figure 1A** (control, galactose and Complex I inhibition). Our moderate-depth global proteome analysis, which was conducted using analytical flow LC, enabled the coverage of 57% of the core mitochondrial ETC complex I – V proteins and 39% of the complex assembly proteins (**Figure 5A**). Analysis of the mitochondrial ETC complex-level protein abundances did not yield any significant differences when comparing the ARID1A WT vs. D9 cells subjected to the various metabolic challenges (**Figure 5B**); however, significant differences were observed at the individual protein level (**Figure 5C**). Loss of ARID1A expression was associated with the significant increase in abundance of four Complex III proteins and eight Complex V proteins, whereas metabolic challenges resulted in the significant increase in the abundance of five Complex I proteins and one Complex IV protein. Results from the pairwise comparisons of the Complex I – V proteins based on ARID1A expression and metabolic challenge are shown in **Supplementary Figures S3-S7.**

**Figure 5.**
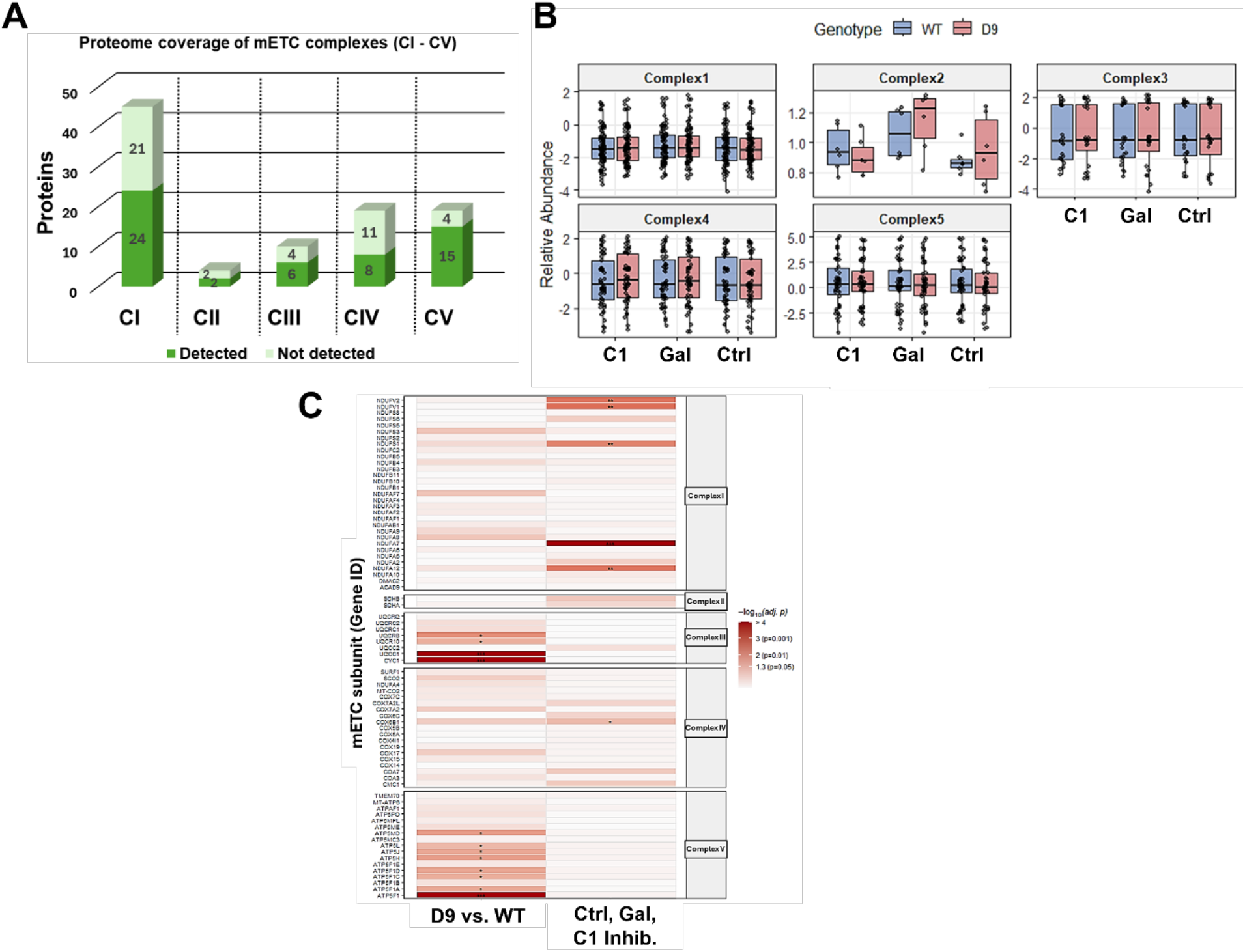
ARID1A knockout is associated with significantly increased abundance of mitochondrial electron transport chain (ETC) complex III and V subunit proteins, whereas metabolic perturbations result in the significantly increased abundance of complex I and IV proteins. **A)** Proteome coverage of mitochondrial ETC complex I – V core proteins. **B)** Relative abundance of mitochondrial ETC complex proteins. Median, inter-quartile range, and individual data points are shown. **C)** Heatmap of statistical significance of mitochondrial ETC complex protein relative abundance differences due to ARID1A loss or metabolic perturbations. Two-way ANOVA with Benjamini-Hochberg FDR correction for multiple comparisons. \**p*.adj<0.05, \*\**p*.adj<0.01, \*\*\**p*.adj<0.001.

To extend the analysis of the mitochondrial ETC complex-level protein abundances (**Figure 5B**), we compared the mean differences (D9 vs. WT) of the proteins in each complex according to the metabolic perturbations (**Figure 6**). Although the differences were not significant (unpaired t-tests with Welch’s correction) (**Figure 6A**), we observed a general trend of increased mitochondrial ETC complex protein abundance in the D9 vs. WT cells (**Figure 6B**). The mean relative abundances of CI, CIII and CIV were higher in the D9 vs. WT cells in the setting of all metabolic challenges, whereas the mean relative abundance of CII in the D9 cells exceeded the abundance in the WT cells only in the control and galactose (Gal) conditions (**Figure 6C**). Treatment with the Complex I (CI) inhibitor was the only condition that resulted in a higher mean relative abundance of the CV proteins in the D9 vs. WT cells.

**Figure 6.**
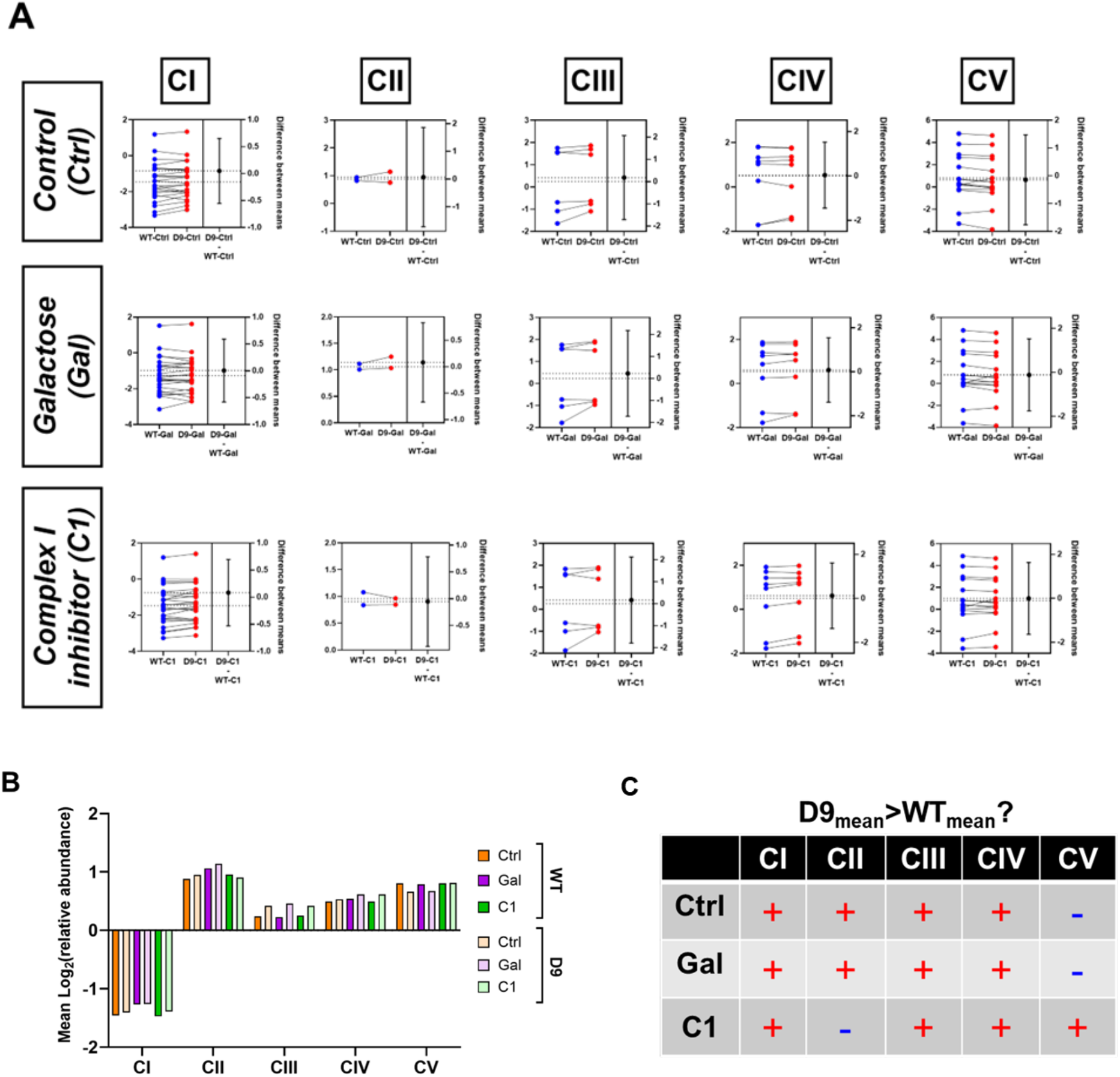
ARID1A knockout and metabolic challenge induce mitochondrial ETC complex-specific perturbations. **A)** Estimation plots of the group mean differences of mitochondrial ETC CI – CV protein relative abundances in the context of ARID1A loss or metabolic challenge. The precision of the calculated effect size is represented as the 95% confidence interval. Dashed horizontal lines indicate the mean of each condition. **B)** Mean relative abundances of CI – CV proteins in the setting of ARID1A expression and metabolic challenge. **C)** Indicators of whether the mean mitochondrial ETC complex protein relative abundances are higher in the absence (D9) vs. presence (WT) of ARID1A expression. **+**: yes, **-**: no.

## DISCUSSION

In this study, we show that ARID1A-deficient RMG1 D9 cells maintain higher mitochondrial respiration than RMG1 WT cells across control, galactose, and Complex I inhibitory conditions. D9 cells displayed higher basal respiration, ATP-linked respiration, maximal respiration, spare respiratory capacity, and coupling efficiency across the conditions examined. These findings are consistent with our previous observations that loss of ARID1A increases mitochondrial respiration and dependence on oxidative metabolism in OCCC cells ^10^. By extending the analysis to conditions that either increase reliance on oxidative phosphorylation through galactose treatment or directly inhibit Complex I, the present study shows that this respiratory phenotype is maintained across distinct metabolic challenges. This is particularly relevant because cancer metabolic phenotypes and dependencies can be strongly influenced by the surrounding metabolic environment. Studies in human tumors and experimental cancer models have shown that nutrient availability and local metabolic conditions can alter nutrient utilization and the response to metabolism-targeting agents ^23–27^.

Another notable observation in this study was that the increased mitochondrial respiration in D9 cells was accompanied by lower steady-state cellular ATP levels. This appears counterintuitive because D9 cells also exhibited increased ATP-linked respiration. However, ATP-linked respiration reflects the portion of oxygen consumption associated with mitochondrial ATP synthesis ^28^, whereas the cellular ATP pool reflects the balance between ATP production and ATP consumption ^28–30^. Thus, increased mitochondrial respiration does not necessarily result in increased steady-state ATP abundance. Increased oxygen consumption without a corresponding increase in ATP production has also been reported in other ARID1A-deficient models ^8^, suggesting that this relationship may represent a broader metabolic feature associated with ARID1A loss.

The basis for the lower ATP levels in D9 cells remains unclear. One possibility is that ARID1A-deficient cells have increased energetic demand and consume ATP at a greater rate, requiring increased mitochondrial respiration to maintain cellular energy requirements. Increased oxygen consumption has been reported alongside increased ATP consumption and lower basal ATP levels in another cellular model ^31^. More recently, mitochondrial ATP consumption has also been shown to support mitochondrial polarization and respiration in acute myeloid leukemia cells, demonstrating that mitochondrial activity can coexist with substantial ATP utilization ^32^. Alternatively, increased respiratory activity may not translate proportionally into ATP production. Mitochondrial uncoupling can produce increased oxygen consumption despite reduced cellular ATP levels ^33,34^, and increased respiration accompanied by reduced ATP synthesis has also been observed under metabolic stress ^35^. However, D9 cells in this study exhibited higher coupling efficiency than WT cells, arguing against a simple model in which the lower ATP abundance is explained primarily by global mitochondrial uncoupling. Together, these observations raise the possibility that increased mitochondrial respiration in ARID1A-deficient cells may reflect increased energetic demand or another underlying metabolic requirement.

Beyond the RMG1 cell model, ARID1A-negative tumors exhibited a higher BEC index than ARID1A-positive tumors in a cohort of 53 human OCCC specimens, suggesting that the association between ARID1A loss and altered mitochondrial bioenergetics is also detectable in clinical disease. A high BEC index was also associated with poorer cancer-specific survival (HR = 5.1, 95% CI 1.4–18; p = 0.0017), suggesting that mitochondrial bioenergetic features may carry prognostic information in OCCC. This is consistent with clinical proteomic analyses in which proteins associated with worse recurrence-free survival were enriched in oxidative phosphorylation-related pathways ^36^. However, the relationship between mitochondrial metabolism and outcome is unlikely to be uniform across OCCC ^37–39^. A proteomic analysis of 192 tumors identified a metabolically active subgroup associated with better progression-free and overall survival after adjustment for clinical variables ^37^. Together, these findings suggest that the clinical significance of mitochondrial metabolism may depend on the specific metabolic program of the tumor rather than simply the overall level of mitochondrial activity.

Proteomic profiling showed that ARID1A status had a greater effect on the RMG1 proteome than the metabolic perturbations. At the individual-protein level, ARID1A loss was associated with significant increases in selected Complex III and Complex V proteins, whereas metabolic challenge was associated with increases in Complex I and Complex IV proteins. Pairwise analysis did not identify significant genotype-dependent differences at the whole-complex level, but the complex-level analysis showed an overall trend toward higher mitochondrial ETC protein abundance in D9 cells. Mean Complex I, III, and IV abundances were higher in D9 across all three conditions, whereas Complex II was higher under control and galactose conditions and Complex V was higher following Complex I inhibition. The overall tendency toward higher ETC complex abundance in D9 cells is consistent with our previous observation that ARID1A loss is associated with increased abundance of mitochondrial ETC proteins ^11^. Consistent with this broader pattern, ARID1A-dependent metabolic remodeling has been reported in other models, including selective changes in mevalonate- and glucose-associated proteins in OCCC, increased oxygen consumption and changes in mitochondrial-associated proteins in ARID1A-deficient cells, and increased OXPHOS protein abundance and TCA-cycle dependence in hepatocellular carcinoma ^8,40,41^. These observations suggest that ARID1A loss is associated with broader metabolic reprogramming, with the specific protein-level changes potentially varying across cellular contexts. Together, our findings indicate that ARID1A loss is associated with ETC protein remodeling, while the increased respiratory phenotype in D9 cells is not accompanied by a uniform increase across all ETC complexes.

These findings further support mitochondrial metabolism as a potential therapeutic vulnerability in ARID1A-deficient OCCC. However, the clinical experience with IACS-010759 highlights the difficulty of directly targeting Complex I, as toxicity limited drug exposure and only modest target inhibition and antitumor activity were achieved at tolerated doses ^42,43^. Continued development of OXPHOS-targeting compounds indicates that mitochondrial metabolism nevertheless remains an active therapeutic area ^20,44–46^. The selective ETC protein changes observed in the present study also underscore the need to define more specific components of the ARID1A-associated mitochondrial dependency rather than relying on broad inhibition of oxidative phosphorylation.

Several limitations should be considered when interpreting these findings. The cell-based functional studies were performed using the RMG1 WT/D9 model, and validation in additional genetically independent ARID1A-deficient models will be important to determine how broadly the observed respiratory phenotype extends across OCCC. In addition, the proteomic analysis measures steady-state protein abundance rather than ETC complex assembly or activity, and coverage of ETC core and assembly proteins was incomplete. Finally, the association between BEC and clinical outcome was evaluated in a single OCCC cohort and will require validation in an independent patient population.

Taken together, our findings show that the ARID1A-associated mitochondrial phenotype persists under distinct metabolic challenges and is not explained simply by coordinated increases in ETC protein abundance. The association of BEC with ARID1A status and cancer-specific survival in human OCCC specimens further links altered mitochondrial bioenergetics with clinical disease. Defining the mechanisms underlying these functional and molecular differences may help identify more specific mitochondrial vulnerabilities in ARID1A-deficient OCCC.

## ACKNOWLEDGEMENTS

The authors thank the Analytical Biochemistry Shared Service of the Masonic Cancer Center for access to CHIMERYS and Proteome Discoverer 3.0. We acknowledge the CRISPR Functional Genomics (CFG) facility at Karolinska Institutet, SciLifeLab, for their support with the generation of CRISPR knockout cell line clones.

## FUNDING

This work was supported by the U.S. Department of Defense Ovarian Cancer Research Program (OC250327P1; M.B.) and the Randy Shaver Community Fund (M.B.). This research was funded in part by the National Cancer Institute, National Institutes of Health, grant R21CA287351 (M.B. and S.N.T.). The content is solely the responsibility of the authors and does not necessarily represent the official views of the National Institutes of Health. This work was supported in part by NIH P30 CA77598 utilizing the following Masonic Cancer Center, University of Minnesota shared resource: Analytical Biochemistry & Mass Spectrometry. This work was supported by the Swedish Cancer Society (Cancerfonden; grant 24 3623 Pj01H; P.D.). B.D. and F.N. were supported by the European Union through PRIN 2022, NextGenerationEU (202237T5AP). B.D. was also partially supported by the Consorzio Interuniversitario Biotecnologie.

## Notes

### Competing Interest Statement

The authors have declared no competing interest.

## REFERENCES

(1) Zhu, C.; Zhu, J.; Qian, L.; Liu, H.; Shen, Z.; Wu, D.; Zhao, W.; Xiao, W.; Zhou, Y. Clinical Characteristics and Prognosis of Ovarian Clear Cell Carcinoma: A 10-Year Retrospective Study. BMC Cancer 2021, 21 (1), 322. 10.1186/s12885-021-08061-7

(2) Rosso, R.; Turinetto, M.; Borella, F.; Chopin, N.; Meeus, P.; Lainè, A.; Ray-Coquard, I.; Le Saux, O.; Ferraioli, D. Ovarian Clear Cell Carcinoma: Open Questions on the Management and Treatment Algorithm. Oncologist 2025, 30 (1), oyae325. 10.1093/oncolo/oyae325

(3) Stewart, J.; Cunningham, N.; Banerjee, S. New Therapies for Clear Cell Ovarian Carcinoma. Int J Gynecol Cancer 2023, 33 (3), 385–393. 10.1136/ijgc-2022-003704

(4) Lebedev, T.; Kousar, R.; Patrick, B.; Usama, M.; Lee, M.-K.; Tan, M.; Li, X.-G. Targeting ARID1A-Deficient Cancers: An Immune-Metabolic Perspective. Cells 2023, 12 (6), 952. 10.3390/cells12060952

(5) Su, Y.-F.; Tsai, E.-M.; Chen, C.-C.; Wu, C.-C.; Er, T.-K. Targeted Sequencing of a Specific Gene Panel Detects a High Frequency of ARID1A and PIK3CA Mutations in Ovarian Clear Cell Carcinoma. Clin Chim Acta 2019, 494, 1–7. 10.1016/j.cca.2019.03.003

(6) Xu, S.; Tang, C. The Role of ARID1A in Tumors: Tumor Initiation or Tumor Suppression? Front Oncol 2021, 11, 745187. 10.3389/fonc.2021.745187

(7) Church, M. C.; Workman, J. L. The SWI/SNF Chromatin Remodeling Complex: A Critical Regulator of Metabolism. Biochem Soc Trans 2024, 52 (3), 1327–1337. 10.1042/BST20231141

(8) Srinivas, U. S.; Tay, N. S. C.; Jaynes, P.; Anbuselvan, A.; Ramachandran, G. K.; Wardyn, J. D.; Hoppe, M. M.; Hoang, P. M.; Peng, Y.; Lim, S.; Lee, M. Y.; Peethala, P. C.; An, O.; Shendre, A.; Tan, B. W. Q.; Jemimah, S.; Lakshmanan, M.; Hu, L.; Jakhar, R.; Sachaphibulkij, K.; Lim, L. H. K.; Pervaiz, S.; Crasta, K.; Yang, H.; Tan, P.; Liang, C.; Ho, L.; Khanchandani, V.; Kappei, D.; Yong, W. P.; Tan, D. S. P.; Bordi, M.; Campello, S.; Tam, W. L.; Frezza, C.; Jeyasekharan, A. D. PLK1 Inhibition Selectively Induces Apoptosis in ARID1A Deficient Cells through Uncoupling of Oxygen Consumption from ATP Production. Oncogene 2022, 41 (13), 1986–2002. 10.1038/s41388-022-02219-8

(9) Wu, S.; Fukumoto, T.; Lin, J.; Nacarelli, T.; Wang, Y.; Ong, D.; Liu, H.; Fatkhutdinov, N.; Zundell, J. A.; Karakashev, S.; Zhou, W.; Schwartz, L. E.; Tang, H.-Y.; Drapkin, R.; Liu, Q.; Huntsman, D. G.; Kossenkov, A. V.; Speicher, D. W.; Schug, Z. T.; Van Dang, C.; Zhang, R. Targeting Glutamine Dependence through GLS1 Inhibition Suppresses ARID1A-Inactivated Clear Cell Ovarian Carcinoma. Nat Cancer 2021, 2 (2), 189–200. 10.1038/s43018-020-00160-x

(10) Zhang, X.; Shetty, M.; Clemente, V.; Linder, S.; Bazzaro, M. Targeting Mitochondrial Metabolism in Clear Cell Carcinoma of the Ovaries. Int J Mol Sci 2021, 22 (9), 4750. 10.3390/ijms22094750

(11) Perez, J. M.; Ryu, J.; Khan, H.; Shetty, M.; Parker, E.; D’Arcy, P.; Zhu, S.; Bazzaro, M.; Thomas, S. N. Proteomic Analysis of ARID1A-Deficient Ovarian Clear Cell Carcinoma Cells Reveals Differential Mitochondria ETC Subunit Abundances and Targetable Mitochondrial Pathways. Int J Mol Sci 2025, 26 (12), 5466. 10.3390/ijms26125466

(12) Elia, I.; Haigis, M. C. Metabolites and the Tumour Microenvironment: From Cellular Mechanisms to Systemic Metabolism. Nat Metab 2021, 3 (1), 21–32. 10.1038/s42255-020-00317-z

(13) Fendt, S.-M.; Frezza, C.; Erez, A. Targeting Metabolic Plasticity and Flexibility Dynamics for Cancer Therapy. Cancer Discov 2020, 10 (12), 1797–1807. 10.1158/2159-8290.CD-20-0844

(14) Stany, M. P.; Vathipadiekal, V.; Ozbun, L.; Stone, R. L.; Mok, S. C.; Xue, H.; Kagami, T.; Wang, Y.; McAlpine, J. N.; Bowtell, D.; Gout, P. W.; Miller, D. M.; Gilks, C. B.; Huntsman, D. G.; Ellard, S. L.; Wang, Y.-Z.; Vivas-Mejia, P.; Lopez-Berestein, G.; Sood, A. K.; Birrer, M. J. Identification of Novel Therapeutic Targets in Microdissected Clear Cell Ovarian Cancers. PLoS One 2011, 6 (7), e21121. 10.1371/journal.pone.0021121

(15) Amano, Y.; Mandai, M.; Yamaguchi, K.; Matsumura, N.; Kharma, B.; Baba, T.; Abiko, K.; Hamanishi, J.; Yoshioka, Y.; Konishi, I. Metabolic Alterations Caused by HNF1β Expression in Ovarian Clear Cell Carcinoma Contribute to Cell Survival. Oncotarget 2015, 6 (28), 26002–26017. 10.18632/oncotarget.4692

(16) Shanmugasundaram, K.; Nayak, B. K.; Friedrichs, W. E.; Kaushik, D.; Rodriguez, R.; Block, K. NOX4 Functions as a Mitochondrial Energetic Sensor Coupling Cancer Metabolic Reprogramming to Drug Resistance. Nat Commun 2017, 8 (1), 997. 10.1038/s41467-017-01106-1

(17) Thomas, L. W.; Esposito, C.; Morgan, R. E.; Price, S.; Young, J.; Williams, S. P.; Maddalena, L. A.; McDermott, U.; Ashcroft, M. Genome-Wide CRISPR/Cas9 Deletion Screen Defines Mitochondrial Gene Essentiality and Identifies Routes for Tumour Cell Viability in Hypoxia. Commun Biol 2021, 4 (1), 615. 10.1038/s42003-021-02098-x

(18) Skinner, O. S.; Blanco-Fernández, J.; Goodman, R. P.; Kawakami, A.; Shen, H.; Kemény, L. V.; Joesch-Cohen, L.; Rees, M. G.; Roth, J. A.; Fisher, D. E.; Mootha, V. K.; Jourdain, A. A. Salvage of Ribose from Uridine or RNA Supports Glycolysis in Nutrient-Limited Conditions. Nat Metab 2023, 5 (5), 765–776. 10.1038/s42255-023-00774-2

(19) Molina, J. R.; Sun, Y.; Protopopova, M.; Gera, S.; Bandi, M.; Bristow, C.; McAfoos, T.; Morlacchi, P.; Ackroyd, J.; Agip, A.-N. A.; Al-Atrash, G.; Asara, J.; Bardenhagen, J.; Carrillo, C. C.; Carroll, C.; Chang, E.; Ciurea, S.; Cross, J. B.; Czako, B.; Deem, A.; Daver, N.; de Groot, J. F.; Dong, J.-W.; Feng, N.; Gao, G.; Gay, J.; Do, M. G.; Greer, J.; Giuliani, V.; Han, J.; Han, L.; Henry, V. K.; Hirst, J.; Huang, S.; Jiang, Y.; Kang, Z.; Khor, T.; Konoplev, S.; Lin, Y.-H.; Liu, G.; Lodi, A.; Lofton, T.; Ma, H.; Mahendra, M.; Matre, P.; Mullinax, R.; Peoples, M.; Petrocchi, A.; Rodriguez-Canale, J.; Serreli, R.; Shi, T.; Smith, M.; Tabe, Y.; Theroff, J.; Tiziani, S.; Xu, Q.; Zhang, Q.; Muller, F.; DePinho, R. A.; Toniatti, C.; Draetta, G. F.; Heffernan, T. P.; Konopleva, M.; Jones, P.; Di Francesco, M. E.; Marszalek, J. R. An Inhibitor of Oxidative Phosphorylation Exploits Cancer Vulnerability. Nat Med 2018, 24 (7), 1036–1046. 10.1038/s41591-018-0052-4

(20) Pujalte-Martin, M.; Belaïd, A.; Bost, S.; Kahi, M.; Peraldi, P.; Rouleau, M.; Mazure, N. M.; Bost, F. Targeting Cancer and Immune Cell Metabolism with the Complex I Inhibitors Metformin and IACS-010759. Mol Oncol 2024, 18 (7), 1719–1738. 10.1002/1878-0261.13583

(21) Cuezva, J. M.; Krajewska, M.; de Heredia, M. L.; Krajewski, S.; Santamaría, G.; Kim, H.; Zapata, J. M.; Marusawa, H.; Chamorro, M.; Reed, J. C. The Bioenergetic Signature of Cancer: A Marker of Tumor Progression. Cancer Res 2002, 62 (22), 6674–6681.

(22) Clemente, V.; Hoshino, A.; Shetty, M.; Nelson, A.; Erickson, B. K.; Baker, R.; Rubin, N.; Khalifa, M.; Weroha, S. J.; Lou, E.; Bazzaro, M. GLS1 Is a Protective Factor in Patients with Ovarian Clear Cell Carcinoma and Its Expression Does Not Correlate with ARID1A-Mutated Tumors. Cancer Res Commun 2022, 2 (8), 784–794. 10.1158/2767-9764.crc-22-0122

(23) Hensley, C. T.; Faubert, B.; Yuan, Q.; Lev-Cohain, N.; Jin, E.; Kim, J.; Jiang, L.; Ko, B.; Skelton, R.; Loudat, L.; Wodzak, M.; Klimko, C.; McMillan, E.; Butt, Y.; Ni, M.; Oliver, D.; Torrealba, J.; Malloy, C. R.; Kernstine, K.; Lenkinski, R. E.; DeBerardinis, R. J. Metabolic Heterogeneity in Human Lung Tumors. Cell 2016, 164 (4), 681–694. 10.1016/j.cell.2015.12.034

(24) Gui, D. Y.; Sullivan, L. B.; Luengo, A.; Hosios, A. M.; Bush, L. N.; Gitego, N.; Davidson, S. M.; Freinkman, E.; Thomas, C. J.; Vander Heiden, M. G. Environment Dictates Dependence on Mitochondrial Complex I for NAD+ and Aspartate Production and Determines Cancer Cell Sensitivity to Metformin. Cell Metab 2016, 24 (5), 716–727. 10.1016/j.cmet.2016.09.006

(25) Abbott, K. L.; Ali, A.; Casalena, D.; Do, B. T.; Ferreira, R.; Cheah, J. H.; Soule, C. K.; Deik, A.; Kunchok, T.; Schmidt, D. R.; Renner, S.; Honeder, S. E.; Wu, M.; Chan, S. H.; Tseyang, T.; Stoltzfus, A. T.; Michel, S. L. J.; Greaves, D.; Hsu, P. P.; Ng, C. W.; Zhang, C. J.; Farsidjani, A.; Kent, J. R.; Madariaga, M. L. L.; Gramatikov, I. M. T.; Matheson, N. J.; Lewis, C. A.; Clish, C. B.; Rees, M. G.; Roth, J. A.; Griner, L. M.; Muir, A.; Auld, D. S.; Vander Heiden, M. G. Screening in Serum-Derived Medium Reveals Differential Response to Compounds Targeting Metabolism. Cell Chem Biol 2023, 30 (9), 1156–1168.e7. 10.1016/j.chembiol.2023.08.007

(26) Rawat, V.; DeLear, P.; Prashanth, P.; Ozgurses, M. E.; Tebeje, A.; Burns, P. A.; Conger, K. O.; Solís, C.; Hasnain, Y.; Novikova, A.; Endress, J. E.; González-Sánchez, P.; Dong, W.; Stephanopoulos, G.; DeNicola, G. M.; Harris, I. S.; Sept, D.; Mason, F. M.; Coloff, J. L. Drug Screening in Human Physiologic Medium Identifies Uric Acid as an Inhibitor of Rigosertib Efficacy. JCI Insight 2024, 9 (13), e174329. 10.1172/jci.insight.174329

(27) Pyun, W. Y.; Park, J. H.; Roh, J. W.; Jeon, D.; Kim, J.; Paik, J. E.; Cho, S. C.; Park, S. Y.; Lim, H.; Kim, H.; Jang, Y. J.; Lee, J.; Byun, J. H.; Son, D. W.; Kim, S.-Y.; Guan, K.-L.; Lee, W. D.; Gee, H. Y.; Lee, H.-W.; No, K. T.; Choi, Y. S.; Hashimoto, T.; Yoshino, T.; Jeong, H.-S.; Namkung, W.; Nam, J. H.; Park, H. W. Nutrient Availability Dictates Cancer Metabolism-Based Therapeutic Responses to Nononcology Drugs. Cancer Res 2026, 86 (5), 1194–1214. 10.1158/0008-5472.CAN-25-1123

(28) Mookerjee, S. A.; Gerencser, A. A.; Nicholls, D. G.; Brand, M. D. Quantifying Intracellular Rates of Glycolytic and Oxidative ATP Production and Consumption Using Extracellular Flux Measurements. J Biol Chem 2017, 292 (17), 7189–7207. 10.1074/jbc.M116.774471

(29) Schmidt, C. A.; Fisher-Wellman, K. H.; Neufer, P. D. From OCR and ECAR to Energy: Perspectives on the Design and Interpretation of Bioenergetics Studies. J Biol Chem 2021, 297 (4), 101140. 10.1016/j.jbc.2021.101140

(30) Desousa, B. R.; Kim, K. K.; Jones, A. E.; Ball, A. B.; Hsieh, W. Y.; Swain, P.; Morrow, D. H.; Brownstein, A. J.; Ferrick, D. A.; Shirihai, O. S.; Neilson, A.; Nathanson, D. A.; Rogers, G. W.; Dranka, B. P.; Murphy, A. N.; Affourtit, C.; Bensinger, S. J.; Stiles, L.; Romero, N.; Divakaruni, A. S. Calculation of ATP Production Rates Using the Seahorse XF Analyzer. EMBO Rep 2023, 24 (10), e56380. 10.15252/embr.202256380

(31) Fang, E. F.; Scheibye-Knudsen, M.; Brace, L. E.; Kassahun, H.; SenGupta, T.; Nilsen, H.; Mitchell, J. R.; Croteau, D. L.; Bohr, V. A. Defective Mitophagy in XPA via PARP-1 Hyperactivation and NAD(+)/SIRT1 Reduction. Cell 2014, 157 (4), 882–896. 10.1016/j.cell.2014.03.026

(32) Hagen, J. T.; Montgomery, M. M.; Aruleba, R. T.; Chrest, B. R.; Krassovskaia, P.; Green, T. D.; Pacheco, E. A.; Kassai, M.; Zeczycki, T. N.; Schmidt, C. A.; Bhowmick, D.; Tan, S.-F.; Feith, D. J.; Chalfant, C. E.; Loughran, T. P.; Liles, D.; Minden, M. D.; Schimmer, A. D.; Shakil, M. S.; McBride, M. J.; Cabot, M. C.; McClung, J. M.; Fisher-Wellman, K. H. Acute Myeloid Leukemia Mitochondria Hydrolyze ATP to Support Oxidative Metabolism and Resist Chemotherapy. Sci Adv 2025, 11 (15), eadu5511. 10.1126/sciadv.adu5511

(33) Papkovskaia, T. D.; Chau, K.-Y.; Inesta-Vaquera, F.; Papkovsky, D. B.; Healy, D. G.; Nishio, K.; Staddon, J.; Duchen, M. R.; Hardy, J.; Schapira, A. H. V.; Cooper, J. M. G 2019S Leucine-Rich Repeat Kinase 2 Causes Uncoupling Protein-Mediated Mitochondrial Depolarization. Hum Mol Genet 2012, 21 (19), 4201–4213. 10.1093/hmg/dds244

(34) Lu, Y.; Li, Y.; Sun, X.; Li, X.; Zhao, S.; Fang, Y.; Qiu, W.; Luo, C.; Qi, W. Mitochondrial Uncoupling Sensitizes Gastric Cancer Cells to Elesclomol-Induced Cuproptosis via FDX1/DLAT Upregulation. Free Radic Biol Med 2026, 244, 284–295. 10.1016/j.freeradbiomed.2025.12.019

(35) Bianchi, G.; Martella, R.; Ravera, S.; Marini, C.; Capitanio, S.; Orengo, A.; Emionite, L.; Lavarello, C.; Amaro, A.; Petretto, A.; Pfeffer, U.; Sambuceti, G.; Pistoia, V.; Raffaghello, L.; Longo, V. D. Fasting Induces Anti-Warburg Effect That Increases Respiration but Reduces ATP-Synthesis to Promote Apoptosis in Colon Cancer Models. Oncotarget 2015, 6 (14), 11806–11819. 10.18632/oncotarget.3688

(36) Yue, L.; Gong, T.; Jiang, W.; Qian, L.; Gong, W.; Sun, Y.; Cai, X.; Xu, H.; Liu, F.; Wang, H.; Li, S.; Zhu, Y.; Zheng, Z.; Wu, Q.; Guo, T. Proteomic Profiling of Ovarian Clear Cell Carcinomas Identifies Prognostic Biomarkers for Chemotherapy. Proteomics 2024, 24 (6), e2300242. 10.1002/pmic.202300242

(37) Ji, J. X.; Cochrane, D. R.; Negri, G. L.; Colborne, S.; Spencer Miko, S. E.; Hoang, L. N.; Farnell, D.; Tessier-Cloutier, B.; Huvila, J.; Thompson, E.; Leung, S.; Chiu, D.; Chow, C.; Ta, M.; Köbel, M.; Feil, L.; Anglesio, M.; Goode, E. L.; Bolton, K.; Morin, G. B.; Huntsman, D. G. The Proteome of Clear Cell Ovarian Carcinoma. J Pathol 2022, 258 (4), 325–338. 10.1002/path.6006

(38) Ji, J. X.; Hoang, L. N.; Cochrane, D. R.; Lum, A.; Senz, J.; Farnell, D.; Tessier-Cloutier, B.; Huntsman, D. G.; Klein Geltink, R. I. The Unique Metabolome of Clear Cell Ovarian Carcinoma. J Pathol 2024, 264 (2), 160–173. 10.1002/path.6329

(39) Xia, S.; Chen, L.; Yu, M.; Li, J.; Chen, J.; Xu, F.; Ni, M.; Liu, C.; Wu, X.; Chen, X.; Li, J. Genetic and Therapeutic Heterogeneity Shape the Baseline and Longitudinal Immune Ecosystem of Ovarian Clear Cell Carcinoma. J Immunother Cancer 2024, 12 (11), e010069. 10.1136/jitc-2024-010069

(40) Goldman, A. R.; Bitler, B. G.; Schug, Z.; Conejo-Garcia, J. R.; Zhang, R.; Speicher, D. W. The Primary Effect on the Proteome of ARID1A-Mutated Ovarian Clear Cell Carcinoma Is Downregulation of the Mevalonate Pathway at the Post-Transcriptional Level. Mol Cell Proteomics 2016, 15 (11), 3348–3360. 10.1074/mcp.M116.062539

(41) Xing, T.; Li, L.; Chen, Y.; Ju, G.; Li, G.; Zhu, X.; Ren, Y.; Zhao, J.; Cheng, Z.; Li, Y.; Xu, D.; Liang, J. Targeting the TCA Cycle through Cuproptosis Confers Synthetic Lethality on ARID1A-Deficient Hepatocellular Carcinoma. Cell Rep Med 2023, 4 (11), 101264. 10.1016/j.xcrm.2023.101264

(42) Yap, T. A.; Daver, N.; Mahendra, M.; Zhang, J.; Kamiya-Matsuoka, C.; Meric-Bernstam, F.; Kantarjian, H. M.; Ravandi, F.; Collins, M. E.; Francesco, M. E. D.; Dumbrava, E. E.; Fu, S.; Gao, S.; Gay, J. P.; Gera, S.; Han, J.; Hong, D. S.; Jabbour, E. J.; Ju, Z.; Karp, D. D.; Lodi, A.; Molina, J. R.; Baran, N.; Naing, A.; Ohanian, M.; Pant, S.; Pemmaraju, N.; Bose, P.; Piha-Paul, S. A.; Rodon, J.; Salguero, C.; Sasaki, K.; Singh, A. K.; Subbiah, V.; Tsimberidou, A. M.; Xu, Q. A.; Yilmaz, M.; Zhang, Q.; Li, Y.; Bristow, C. A.; Bhattacharjee, M. B.; Tiziani, S.; Heffernan, T. P.; Vellano, C. P.; Jones, P.; Heijnen, C. J.; Kavelaars, A.; Marszalek, J. R.; Konopleva, M. Complex I Inhibitor of Oxidative Phosphorylation in Advanced Solid Tumors and Acute Myeloid Leukemia: Phase I Trials. Nat Med 2023, 29 (1), 115–126. 10.1038/s41591-022-02103-8

(43) Machado, N. D.; Heather, L. C.; Harris, A. L.; Higgins, G. S. Targeting Mitochondrial Oxidative Phosphorylation: Lessons, Advantages, and Opportunities. Br J Cancer 2023, 129 (6), 897–899. 10.1038/s41416-023-02394-9

(44) Cao, L.; Yao, H.; Hou, P.; Ren, Y.; Zhi, H.; Jia, X.; Li, X. Synthesis and Evaluation of New 5-(1H-1,2,4-Triazol-3-Yl)-1,2,4-Oxadiazole Derivatives and Their Application as OXPHOS Inhibitors. Eur J Med Chem 2025, 297, 117903. 10.1016/j.ejmech.2025.117903

(45) Vasan, K.; Chandel, N. S. Molecular and Cellular Mechanisms Underlying the Failure of Mitochondrial Metabolism Drugs in Cancer Clinical Trials. J Clin Invest 2024, 134 (3), e176736. 10.1172/JCI176736

(46) Beerkens, A. P. M.; Heskamp, S.; Reinema, F. V.; Adema, G. J.; Span, P. N.; Bussink, J. Mitochondria Targeting of Oxidative Phosphorylation Inhibitors to Alleviate Hypoxia and Enhance Anticancer Treatment Efficacy. Clin Cancer Res 2025, 31 (7), 1186–1193. 10.1158/1078-0432.CCR-24-3296

