## Supplemental Figure 1-7 for "Bioenergetic and Proteomic Profiling of ARID1A-Deficient Ovarian Clear Cell Carcinoma under Metabolic Challenge"

#### Supporting Information

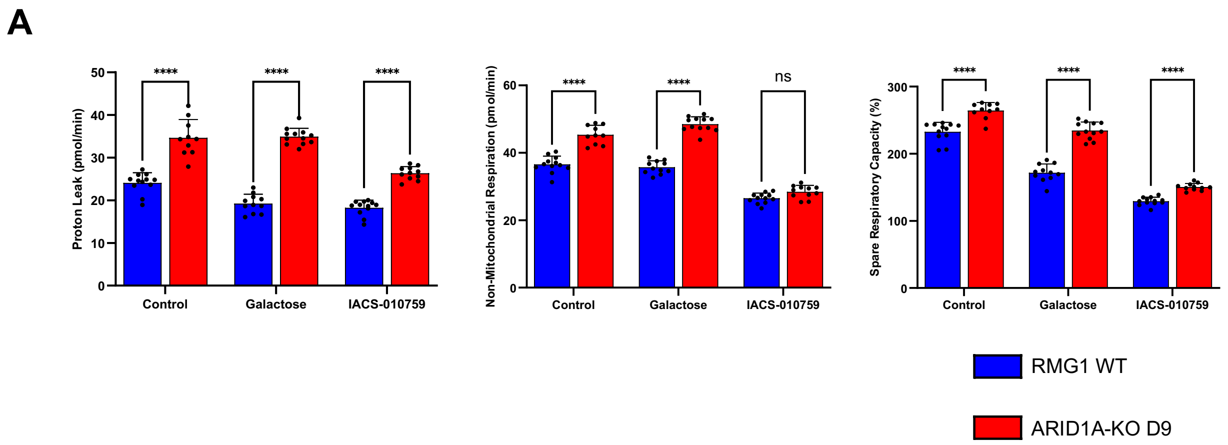


**Figure S1. Additional mitochondrial respiratory parameters in RMG1 WT and D9 cells.** (A) Proton leak, non-mitochondrial respiration, and spare respiratory capacity expressed as a percentage of basal respiration in RMG1 WT and ARID1A-knockout D9 cells under the indicated metabolic conditions. Data points represent individual Seahorse wells and bars represent mean ± SEM.

| Sample | Range of biological replicate protein abundance correlations (R) |
|---|---|
| WT Ctrl | 0.992 – 0.993 |
| D9 Ctrl | 0.994 – 0.996 |
| WT Gal | 0.994 – 0.995 |
| D9 Gal | 0.987 – 0.991 |
| WT C1 | 0.989 – 0.994 |
| D9 C1 | 0.993 – 0.995 |

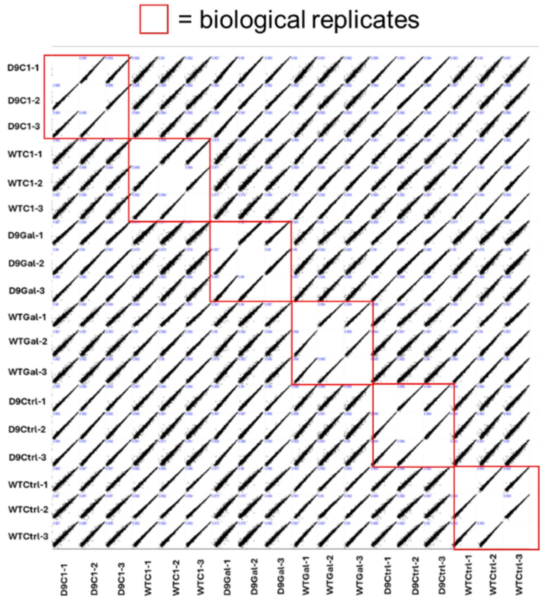


**Figure S2. Correlation plots of protein abundances across all samples.** R = Pearson correlation. n = 3 biological replicates per condition.

### Complex I proteins

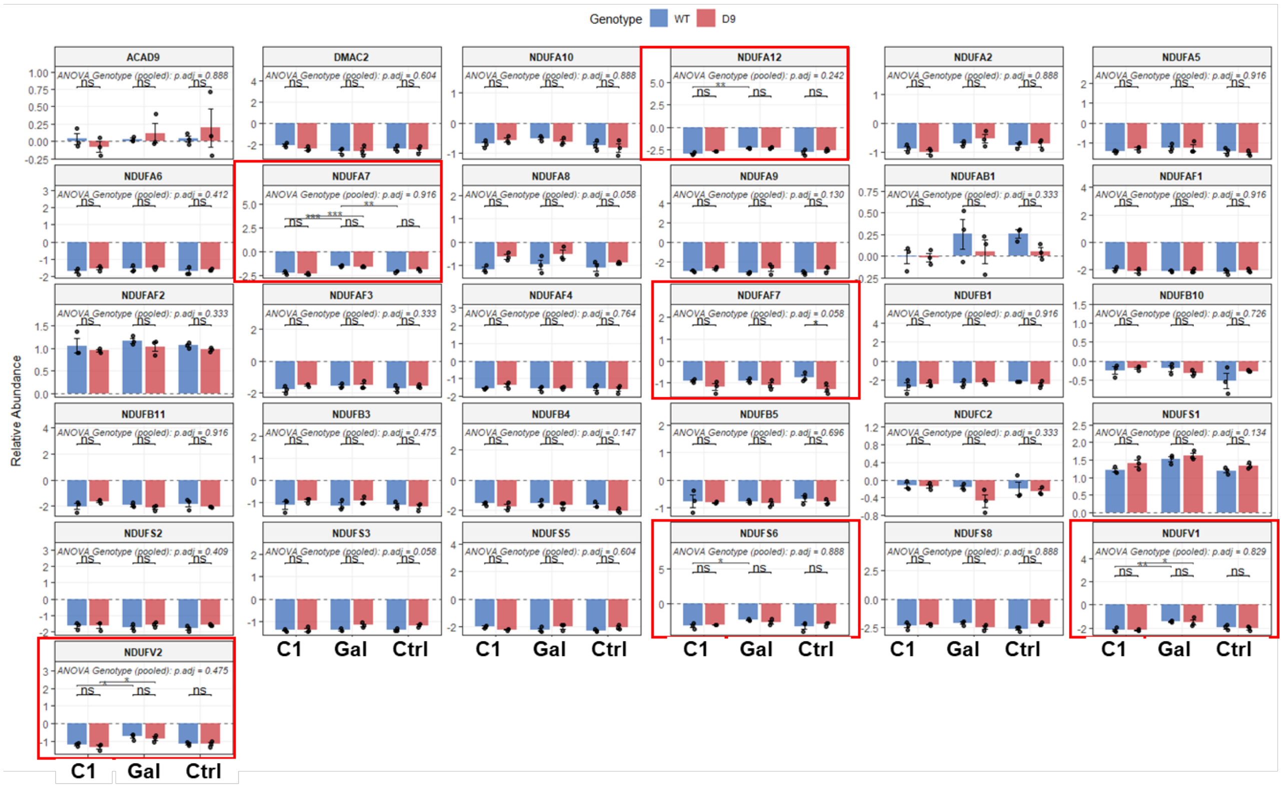


**Figure S3. Pairwise comparisons of Complex I protein abundances.** Red boxes indicate proteins with significantly different protein abundances between genotypes or metabolic conditions. Limma-moderated t-tests with Benjamini Hochberg-adjusted FDR. *$p$.adj<0.05, **$p$.adj<0.01, ***$p$.adj<0.001.

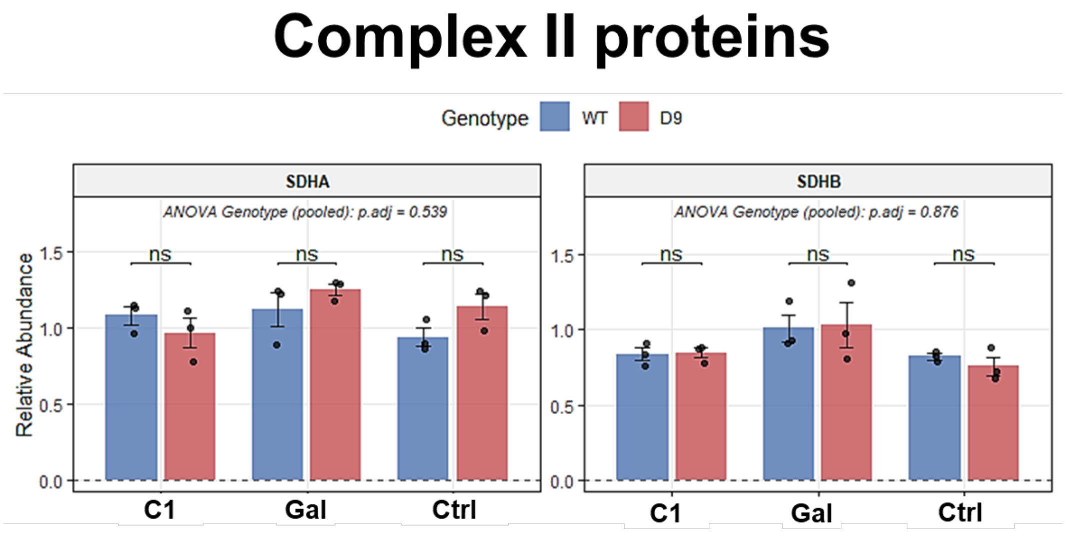


**Figure S4. Pairwise comparisons of Complex II protein abundances.** Limma-moderated t-tests with Benjamini Hochberg-adjusted FDR. **p*.adj<0.05, ***p*.adj<0.01, ****p*.adj<0.001.

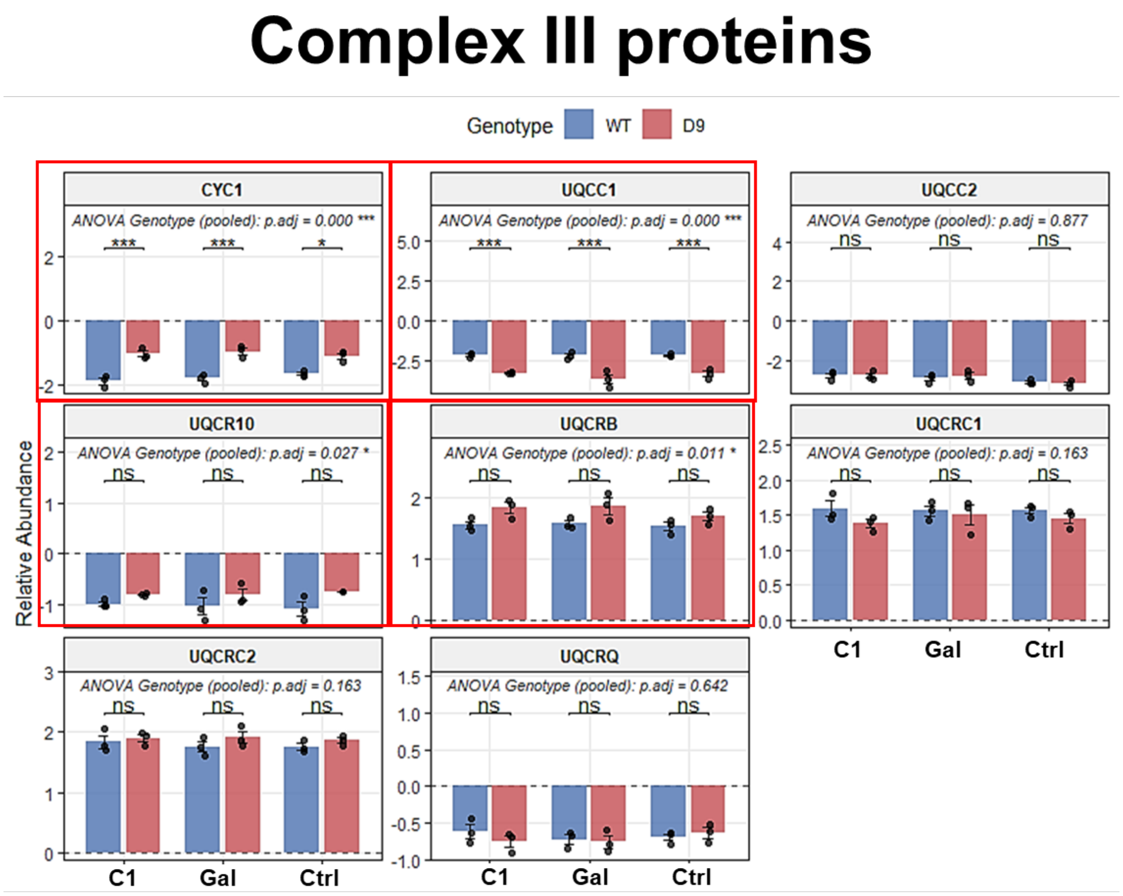


**Figure S5. Pairwise comparisons of Complex III protein abundances.** Red boxes indicate proteins with significantly different protein abundances between genotypes or metabolic conditions. Limma-moderated t-tests with Benjamini Hochberg-adjusted FDR. **p*.adj<0.05, ***p*.adj<0.01, ****p*.adj<0.001.

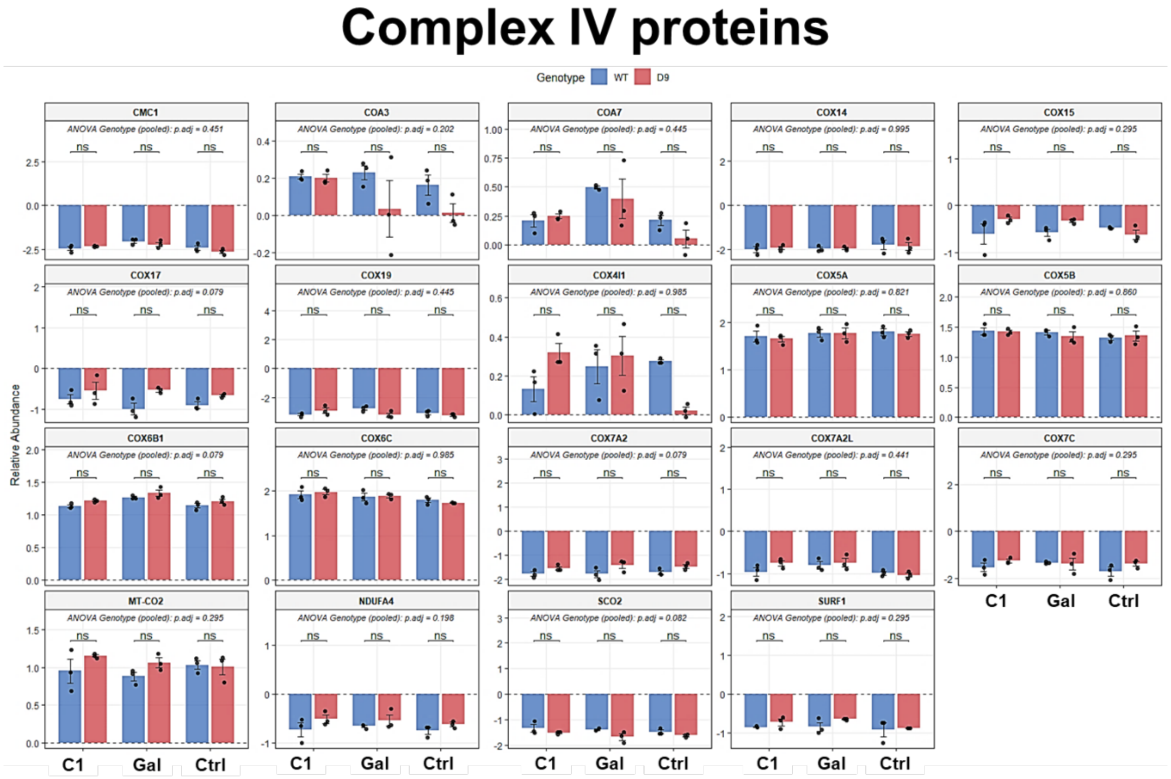


**Figure S6. Pairwise comparisons of Complex IV protein abundances.** Limma-moderated t-tests with Benjamini Hochberg-adjusted FDR. **p*.adj<0.05, ***p*.adj<0.01, ****p*.adj<0.001.

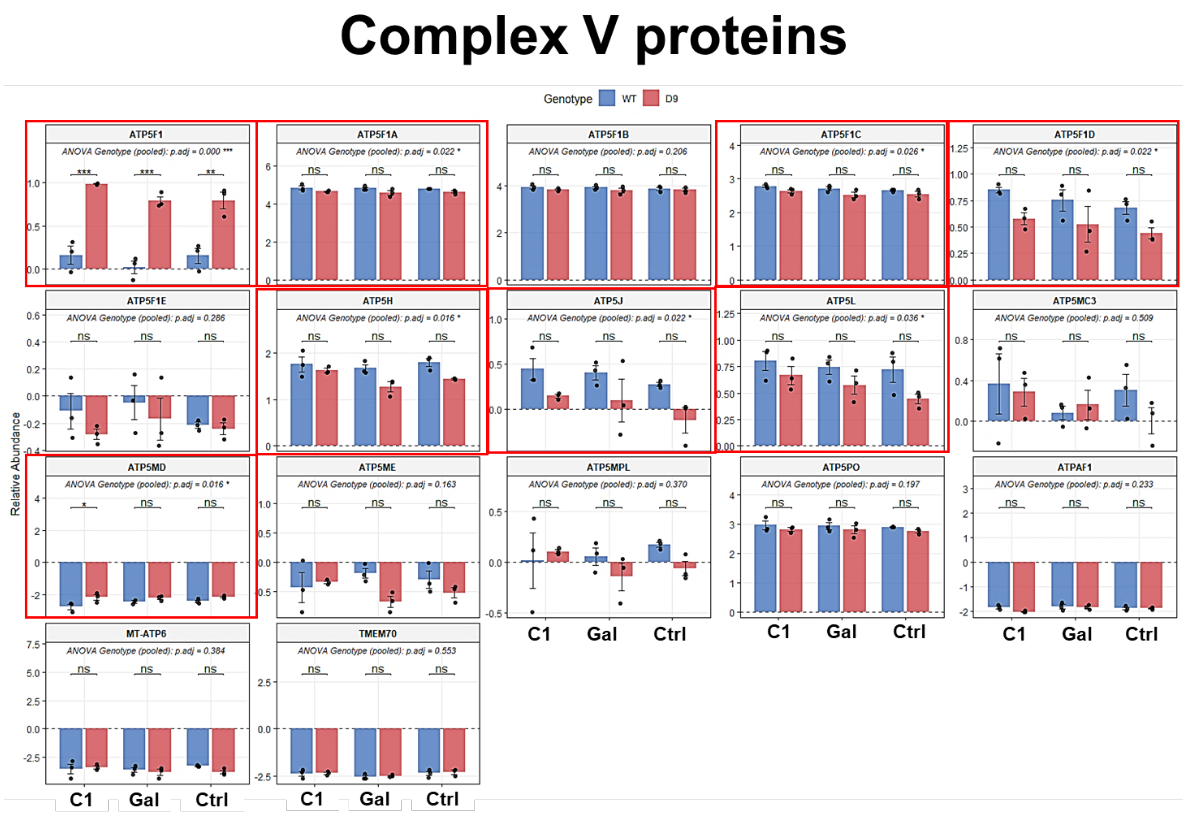


**Figure S7. Pairwise comparisons of Complex V protein abundances.** Red boxes indicate proteins with significantly different protein abundances between genotypes or metabolic conditions. Limma-moderated t-tests with Benjamini Hochberg-adjusted FDR. *$p$.adj<0.05, **$p$.adj<0.01, ***$p$.adj<0.001.
